# Identifying genomic surveillance gaps in Africa for the global public health response to West Nile Virus

**DOI:** 10.1101/2024.12.18.629123

**Authors:** Monika Moir, Nikita Sitharam, Marije Hofstra, Graeme Dor, Gaspary Mwanyika, Yajna Ramphal, Martina L. Reichmuth, San Emmanuel James, Robert Gifford, Eduan Wilkinson, Derek Tshiabuila, Wolfgang Preiser, Abla Ahouefa Konou, Molalegne Bitew, Bernard Anyebe Onoja, Giacomo Maria Paganotti, Adugna Abera, James Ayei Maror, John Kayiwa, Sara Abuelmaali, Eddy Kinganda Lusamaki, CLIMADE Consortium, Marietjie Venter, Felicity Burt, Cheryl Baxter, Richard Lessells, Tulio de Oliveira, Houriiyah Tegally

## Abstract

**Background:** West Nile Virus (WNV) is a zoonotic flavivirus of significant One Health relevance and is classified as a priority pathogen with a high-risk of causing public health emergencies of global concern. WNV is endemic to Africa; however, the availability of genomic sequences from the continent remains limited.

**Methods:** We review the extent of polymerase chain reaction testing and genomic sequencing of WNV conducted across Africa. Using phylogeographic methods, we map the spatiotemporal spread of the virus across the continent and globally.

**Findings:** Our study shows that WNV has been detected in 39 African countries (including Comoros, Seychelles, and Mauritius), the Canary Islands, and Réunion Island. Publications including molecular data originate from 24 countries; however, genomic sequences are publicly available for only 16 countries. We identify regions with detected viral circulation but lacking molecular surveillance. Further, we list such regions that overlap with Key Biodiversity Areas (sites harbouring significant bird diversity) as they may host high viral circulation, and high human population density that may be susceptible to spillover.

**Interpretation:** We recognise significant knowledge gaps on the true disease burden, molecular epidemiology, and distribution of WNV in Africa. Addressing these gaps requires an integrated One Health surveillance approach which is challenging to establish. We propose three key surveillance needs as potential starting points to improve our understanding of the virus in Africa to strengthen the global public health response to this disease.

**Funding:** Rockefeller Foundation, the National Institute of Health USA, Institute of Human Virology Nigeria, Global Health EDCTP3 Joint Undertaking, the Health Emergency Preparedness and Response Umbrella Program, managed by the World Bank Group, the Medical Research Foundation, and the Wellcome Trust.

## Introduction

West Nile Virus (WNV) is a zoonotic flavivirus that is maintained in a transmission cycle between birds and vector-competent mosquitoes, but may be transmitted incidentally to dead-end hosts such as humans, horses, other mammal species (1), and reptiles (2). WNV was first isolated in 1937 from a febrile patient in the West Nile district of Uganda, as a neurotropic virus causing lesions to the central nervous system (3). Although most WNV infections are clinically silent, symptoms and severity vary between infected hosts. In humans, 80% of infections are asymptomatic, and symptomatic infections typically present as a mild febrile illness with headaches, myalgia, arthralgia and rash. Neuroinvasive diseases such as encephalitis or meningitis are observed in less than 1% of cases but have a fatality rate of 10-30% (4). Approximately 20% of infected horses develop clinical signs, 90% of which display neurological symptoms of limb ataxia, paralysis, and seizures (5). Natural avian infections may also present with neurological signs similar to that seen in humans, inflammation, and necrosis of major organ systems, bone marrow, and skeletal muscles have also been found (6).

WNV is of global public health significance and has been included on the World Health Organisation (WHO) list of priority pathogens with a high risk of causing public health emergencies of international concern for all global regions (7). It has caused morbidity and mortality in humans for several decades. For example, one of the largest epidemics occurred in the Karoo region of South Africa in 1974 causing tens of thousands of cases (8), and Tunisia has experienced recurring outbreaks in 1997, 2003, 2012, and 2018 (9). Over the last two decades, WNV emerged as an important public health concern in the Americas and Europe, causing over 25,000 cases of neurological diseases in the first decade after emergence in the USA (10). Also, an increasing number of affected regions in Europe indicate an expanding geographical range of WNV (11). As such, WNV occupies a distinct position within the landscape of arboviral diseases as it currently has an almost global distribution and has caused one of the largest outbreaks of neuroinvasive disease in humans (12), yet its transmission patterns remain one of the least understood.

WNV is a disease of One Health importance and is transmitted primarily by mosquitoes of the *Culex* genus. In Africa, it has been isolated from 46 mosquito species of seven genera with the vectorial competence of 38 species still needing assessment (13). The most important WNV vectors are members of the *Culex pipiens* species complex as they exhibit high viral transmission rates (14), and *Cx. univittatus* due to its ornithophilic feeding behaviour (15) and the proven vertical transmission of WNV in this species (16). *Cx. quinquefasciatus* may be a major vector in urban settings due to its distribution in urban areas and activity through all seasons of the year (17). WNV has also been detected in ticks such as *Rhipicephalus pulchellus* sampled from livestock in Kenya (18). Ticks may spread WNV over long distances via birds but their role in transmission of the virus requires further research.

WNV was first detected in Africa and is endemic to many regions of the continent (19). Despite this, the known distribution, transmission, and epidemiological patterns on the continent are patchy (13). Numerous African countries have experienced a long-term burden of infections but there remains a gap in identifying the true distribution and transmission risk in Africa, with epidemiological knowledge lacking for 19 countries on the continent (13). Genomic data is vital for investigating transmission dynamics and understanding disease outbreaks (20–22), but publicly available African WNV sequences are mostly limited to gene fragments with sparse geographical representation. This is partly due to the low viremia in humans or horses with clinical signs that have a short molecular diagnostic window and most cases being diagnosed through serology. Nevertheless, considering the power of such genomic epidemiological analyses, a full assessment of the landscape of WNV molecular surveillance is needed. Therefore, we conducted a review of publications to assess the extent of polymerase chain reaction (PCR) testing and genomic sequencing that has been performed on WNV across Africa. We mapped the occurrence of the virus and molecular studies for different host species and vectors to identify areas in which WNV circulation is confirmed but molecular data is absent. We highlight regions where molecular investigations would be beneficial.

Also, considering the widespread distribution of WNV and the severity of outbreaks in naive populations, it is important to understand the global historical dispersal from its inferred African origin (13,20). Recent studies present phylogeographic reconstructions limited to certain regions of the world (23–25). Here, we use publicly available whole genomes to reconstruct the spatiotemporal spread of the virus within the African continent and globally.

## Methods

### Literature review of WNV genomic sequencing and PCR testing in Africa

To assess the landscape of molecular surveillance (PCR testing and genomic sequencing) of WNV in Africa, we undertook a systematic review according to Preferred Reporting Items for Systematic reviews and Meta-Analyses (PRISMA) criteria (26). The review protocol was registered with the International prospective register of systematic reviews (PROSPERO ID no. CRD42024614647). Articles were obtained with search terms pertaining to the virus and disease name, geographic locations in Africa, and genomic or nucleic acid terms from PubMed (n=248) and Scopus (n=263) databases on 29 April 2024 (Supplementary information S1). We did not restrict the date or language of publication. We limited our search to published peer-reviewed articles and found additional publications through searches of references in retrieved articles (n=3).

We removed duplicates and two reviewers independently screened the remaining articles (n=334) by title and abstract. We selected publications for full-text screening if they satisfied the following criteria: i) WNV infections were natural (excluded experimental infections), ii) Produced or analysed nucleic acid data via PCR or genomic sequencing methods (publications were excluded if they employed other diagnostic techniques but were included if they used a combination of techniques that included molecular diagnostics), iii) Molecular data was produced from samples that were collected in Africa. We included publications that produced or analysed molecular data from islands that are geographically close to the African continent but not politically classed as African nations (for example Réunion and Canary Islands) as birds move between continental Africa and many of these islands (27,28) with the potential for WNV transmission opportunities between these areas. Two reviewers extracted relevant data (and checked the accuracy of each other’s extractions) from the included publications (n=79) (Supplementary information S2). Extracted data included study design, date of sampling collection and fine-scale location information, molecular methods used, host and sample type, positive and negative detections of WNV, and sequence publication information. We classed each study as to whether it had a focus on Africa or an African country, if the focus of the study was elsewhere but analysed sequences from Africa, or the study focus was elsewhere but it produced and/or analysed sequences from Africa.

Two reviewers categorised (and checked one another’s categorisation) the publications into a study type with the following definitions: *Indicator-based surveillance* - from the Technical Guidelines for Integrated Disease Surveillance and response of the World Health Organization (WHO) African Region (29), the regular identification, collection, monitoring, analysis, and interpretation of data produced by formal sources such as facility-based surveillance, sentinel surveillance, syndromic surveillance. *Event-based surveillance* - the quick capture of information following events that are of potential risk to public health such as unusual disease or deaths, atypical clustering of cases, events in the community or environment causing potential exposure to disease (29). *Case study* - clinical case reports of patients with classical or unusual presentations of the disease and studies describing the episode of diagnosis and care for a patient (30). *Travel-related surveillance* - a case of a traveller either entering an African country or departing from an African country; or regional movement between African countries. *Research* - studies of the transmission, characterisation, diversity, etc. of WNV that are not done as part of the above-listed surveillance strategies.

To determine whether our review captured all available sequences, we cross-referenced the sequence information we extracted from publications (n=257 sequences) with those that are publicly available on NCBI (n=289) (Supplementary information S3). Of the sequences not captured by our review, we found 40 were not linked to publications, and 18 were published but did not meet our criterion of being sampled from natural infections. We identified 26 sequences from our review that are not publicly accessible.

### Records of WNV detection from reported cases, deaths, seroprevalence surveys, and other research studies

In order to compare where WNV molecular studies have been conducted relative to all locations that the virus has been recorded (from reported cases, seroprevalence surveys, and other research studies in all host and vector species in Africa), we used the Global Infectious Diseases and Epidemiology Network (GIDEON) West Nile Fever: Global Status report 2023 (31) as a guide for available literature. We reviewed all primary sources referenced in the GIDEON report to collect records of WNV with as fine-scale location data as possible. From these sources, we also extracted the year of study, host species, number of reported cases/deaths, percentage positivity from seroprevalence surveys, and tests performed to ascertain WNV positivity. We also performed further nonsystematic searches for additional reports of WNV from published papers, reviews, and ProMed alerts to find sources not listed in the GIDEON report. When mapping the total number of viral occurrences relative to molecular sampling locations, all WNV records were flattened to one occurrence per sampling location point (irrespective of the number of cases or seroprevalence results). The same was done for the molecular studies.

We produced data visualisations with RStudio Version 2024.04.2+764 (32) and QGIS version 3.26.3 (33). If detailed location data was not available for plotting the maps, points were positioned at the centroid of the country of collection. This was done for at least one study for Algeria, Egypt, Central African Republic, Ivory Coast, Kenya, Madagascar, Nigeria, Senegal, South Africa, Tunisia, Uganda and Zambia.

### Phylogenetic analyses

We retrieved whole genome sequences from NCBI Virus (https://www.ncbi.nlm.nih.gov/labs/virus/vssi/#/) on 13 August 2024 with a filter of 10 kilobase (kb) minimum sequence length. We retrieved sequences from all host and vector species. We subsampled sequences from the United States of America with the NCBI Virus function of randomly selecting three sequences per sample collection year (n = 66). We manually screened sequences for quality to confirm <10% base ambiguity and removed sequences lacking collection date and sampling locations. We analysed sequences through Genome Detective West Nile Virus Typing Tool for lineage assignment (34). We selected only Lineage 1A (L1A) and Lineage 2 (L2) sequences for further analysis as these are the most prevalent lineages in Africa (19). Lineage specific alignments were performed in MAFFTv7 (35). We utilised RDP5.59 to perform a full recombination screening with the two alignments to identify recombinant sequences (36). We identified six recombinants that were removed from alignments for further analyses (OP734266, OP846974, MF797870 for L1A and OK239667, EF429199, OQ214888 for L2). Additionally, we removed sequences of highly passaged isolates or clonal variants. The final alignments contained 258 and 541 sequences for Lineage 1A and Lineage 2, respectively.

We reconstructed phylogenetic trees and phylogeographic dispersal separately for L1A and L2 whole genome alignments. We constructed maximum likelihood phylogenies with IQTree v2.3.6 (37) including a search for best-fitting models with ModelFinder Plus (best-fit model for L1A was GTR+F+I+R3 according to Bayesian Information Criterion; and GTR+F+I+R4 for L2) and 1,000 bootstrap approximations. We inspected the tree topologies for temporal molecular clock signals using the *clock* functionality of TreeTime (38), removed outlier sequences (n=2 for L1A and n=8 for L2) that deviated from the strict molecular clock assumption, and created time-scaled phylogenies using adjusted mutation rates for each lineage (lJ=4.35 ✕ 10^-4^ for L1A and lJ=2.62 lJ 10^-4^ for L2) as determined in a root-to-tip regression analysis (Supplementary information S4 and S5). We ran the *mugration* model with the time-scaled phylogenies in TreeTime to annotate discrete country locations to tips and infer country locations for internal nodes. Geographical resolution was at the country level for all sequences. Using a custom python script, we quantified the state changes over the phylogenies from the roots toward the tips. Visualisation and annotation of phylogenetic trees and viral exchange mapping were done in R. The maps link the origin to the destination country with curved lines going anti-clockwise in the direction of the curve. Each curved line is coloured by the mean date of state changes occurring along that specific link.

## Results

### African WNV molecular surveillance is mostly driven by research

We identified 79 publications employing molecular methods to study WNV in Africa since 1990, with a gradual increase in publication rate to a peak in 2021 (Figure 1A). The earliest publication isolated WNV strains from hepatitis patients in Bangui, Central African Republic (CAR), flagging for the first time the possibility of liver involvement (39). Of the 79 publications, 56 had an African focus and 23 had a primary study focus outside of Africa. Of those 23, three produced African genomic data and 20 employed African sequences in their analyses (Figure 1A). The three studies that produced African genomic data sequenced stored samples primarily for performing phylogenetic and phylogeographic analyses for regions of interest other than Africa. Of the 56 studies with an African focus, 18 used human samples, 15 animal hosts, 15 vector samples, and eight publications screened multiple host or vector groups (Figure 1A).

**Figure 1:**
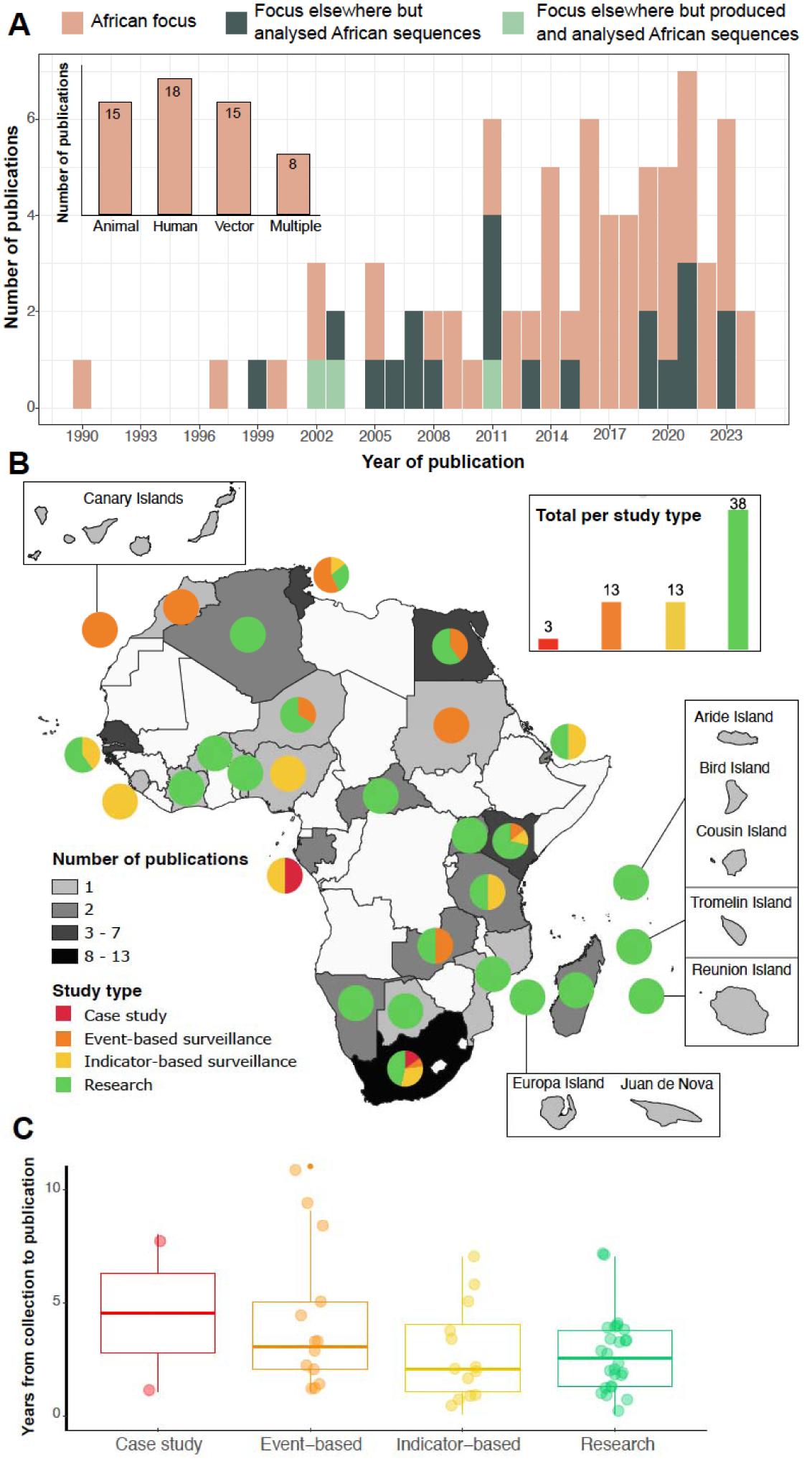
Publications of West Nile Virus genomics in Africa and distribution of study types. A) Annual number of publications utilising molecular methods to study West Nile Virus in Africa. Studies are categorised based on whether the geographic focus is Africa or elsewhere in the world and if molecular data is produced and/or analysed from African samples. The inset bar graph depicts, for studies with an African focus, the total number of published studies per host or vector group, where ‘Multiple’ indicates a combination of at least two different host or vector groups; B) Publications with an African focus were mapped with pie charts to show the proportions of study type categories for each country, while the colour ramp displays the total number of studies per country. The inset bar graph depicts the total publications per study type. Islands for which we found published studies are zoomed in peripheral boxes; C) Box plots show the distribution of publication turnaround time (measured as the number of years from sample collection to study publication) for each study type. Individual data points are displayed by circles. One outlier with a 25-year turnaround time from the research study type was removed for improved visual resolution of the differences between study types.

Considering only the studies with an African focus, we retrieved publications for 24 countries of continental Africa, with the most publications originating from South Africa, followed by Senegal, Tunisia, Egypt, and Kenya (Figure 1B). The vast majority were research studies (n=38) which were conducted across most of the continent (Figure 1B). Case studies (n=3) were published only from Gabon and South Africa. Event-based surveillance studies (n=13) were published from eight countries of continental Africa as well as the Canary Islands, while indicator-based surveillance studies (n=13) were published from nine different countries. Indicator-based surveillance studies and event-based surveillance showed an average turnaround time from sample collection to study publication, of 2.7 and 4.1 years, respectively (Figure 1C). Research publications and case study analyses had an average turnaround time of 3.5 and 4.5 years, respectively. No travel-related surveillance studies were identified. For 31 of 55 African countries, we did not identify molecular publications on WNV, as shown by the unshaded areas in Figure 1B.

### Molecular testing for WNV infection in humans remains limited

In our review of WNV molecular papers, we extracted and collated PCR/sequencing confirmed WNV infections, which are indicative of active infections, for human cases (Table 1), animal hosts (Table 2), and vectors (Table 3).

**Table 1:**
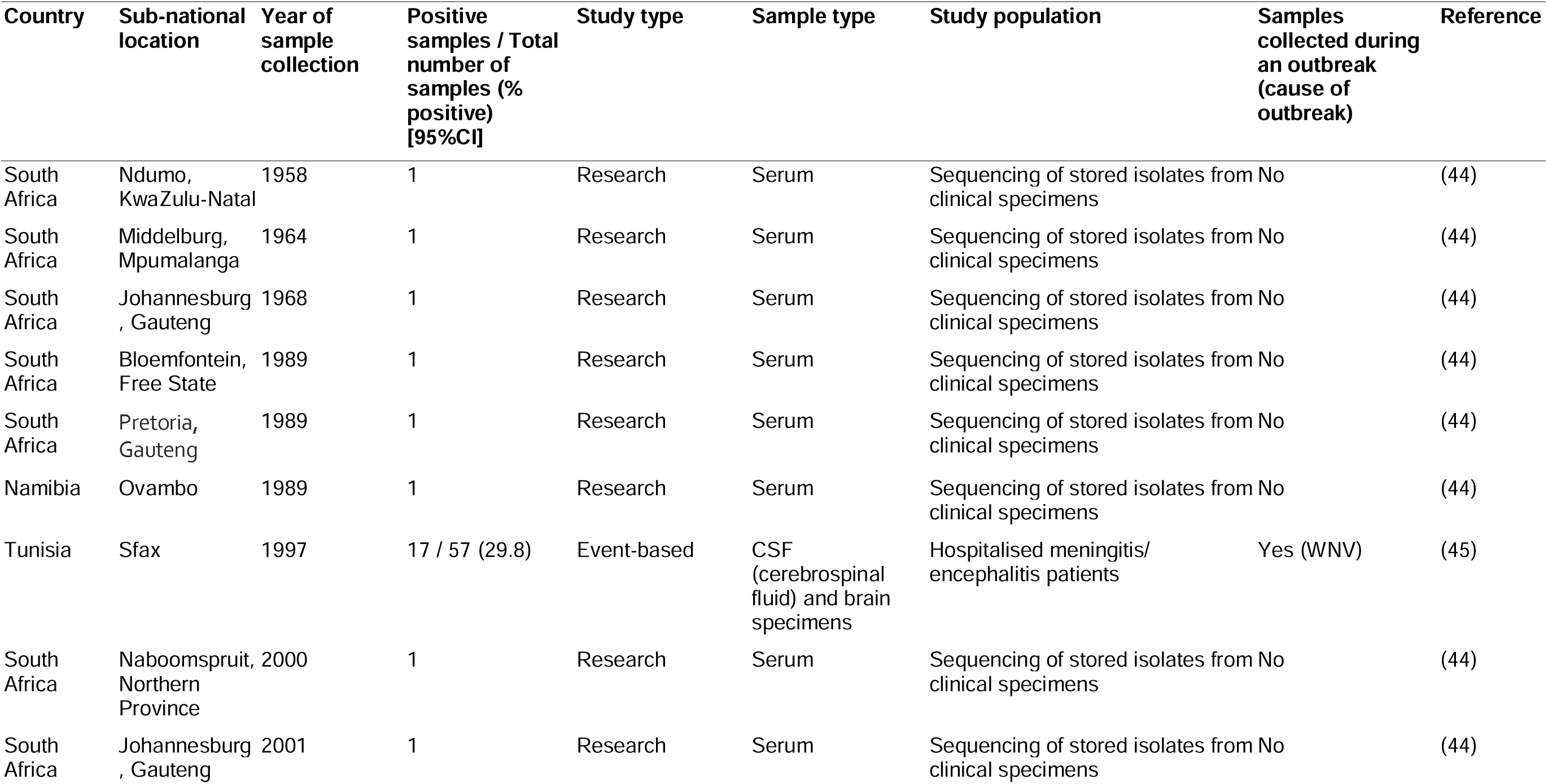
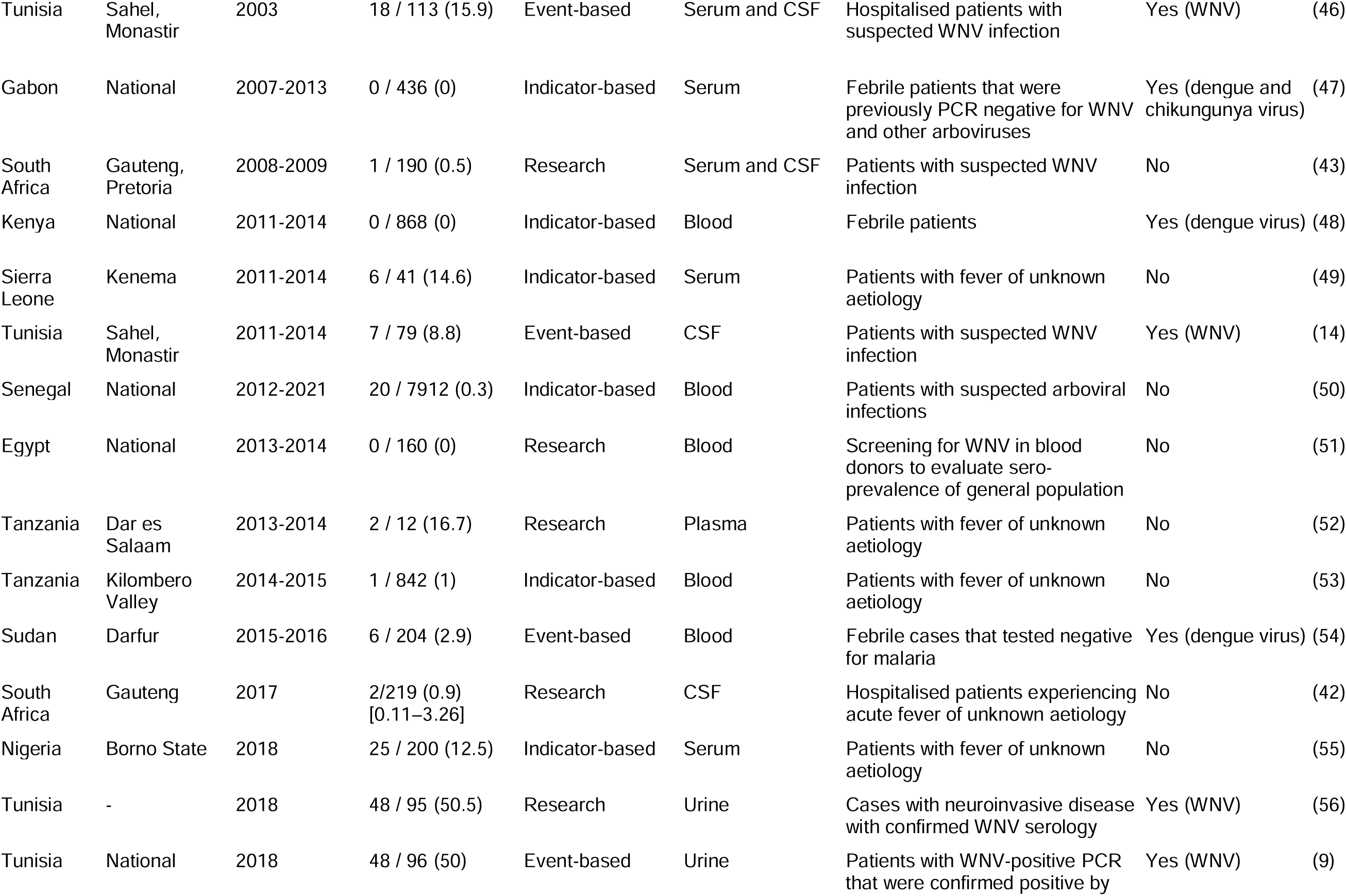
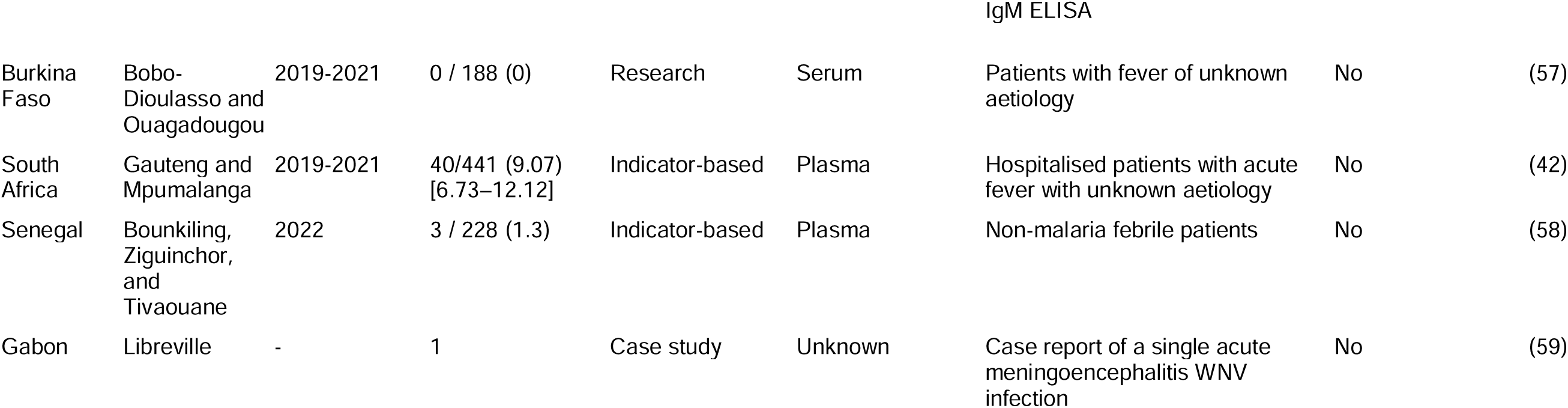
PCR testing and genomic sequencing of West Nile Virus in humans across Africa. Contextual information of sample collection and WNV testing results are listed. Study type categorisation of indicator-based surveillance, event-based surveillance, research or case study is listed. Sub-national location information is blank for studies from which sample collection location could not be retrieved. 95% confidence intervals (CI) of sample positivity are provided if available from the primary study.

**Table 2:**
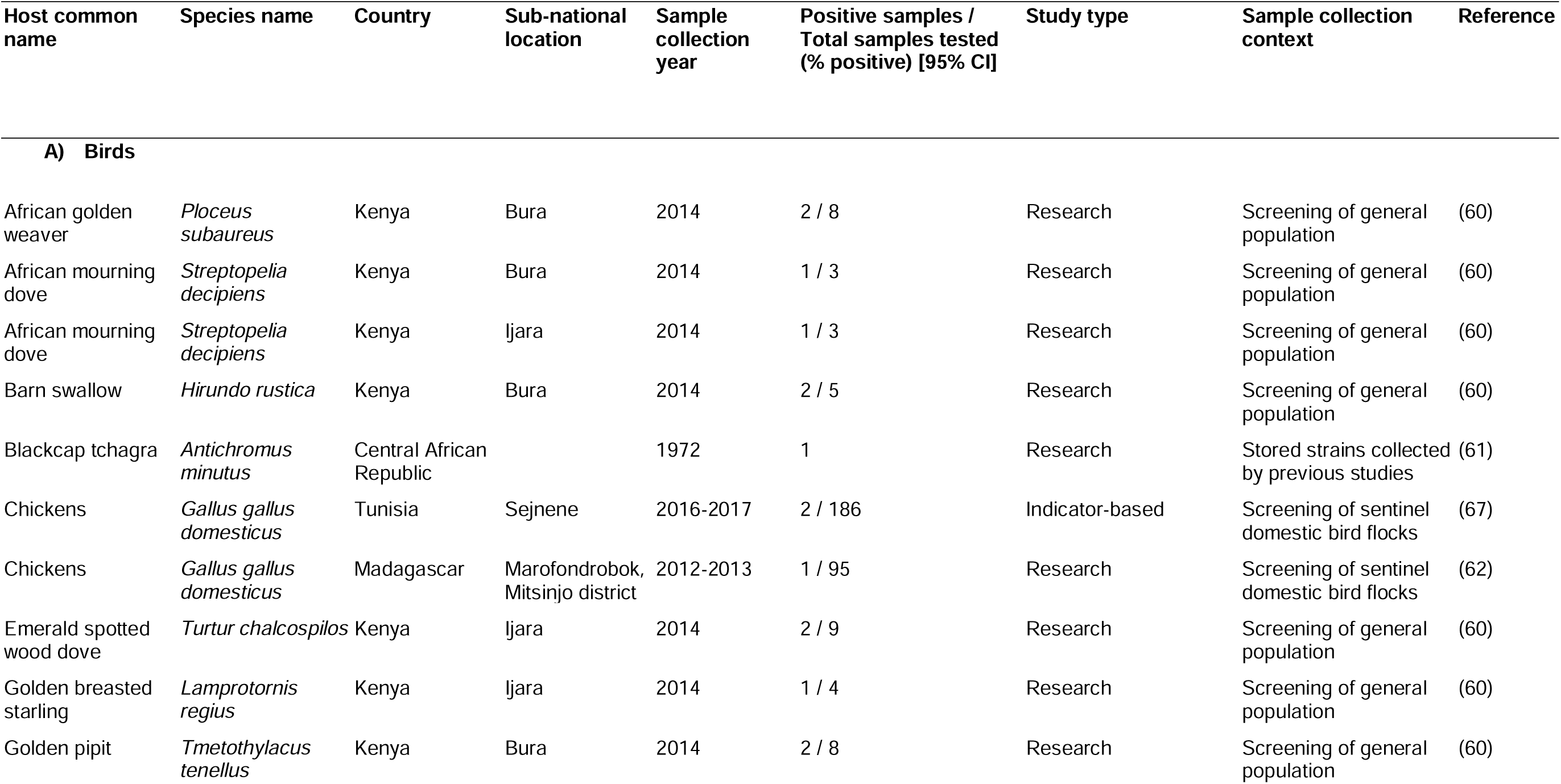
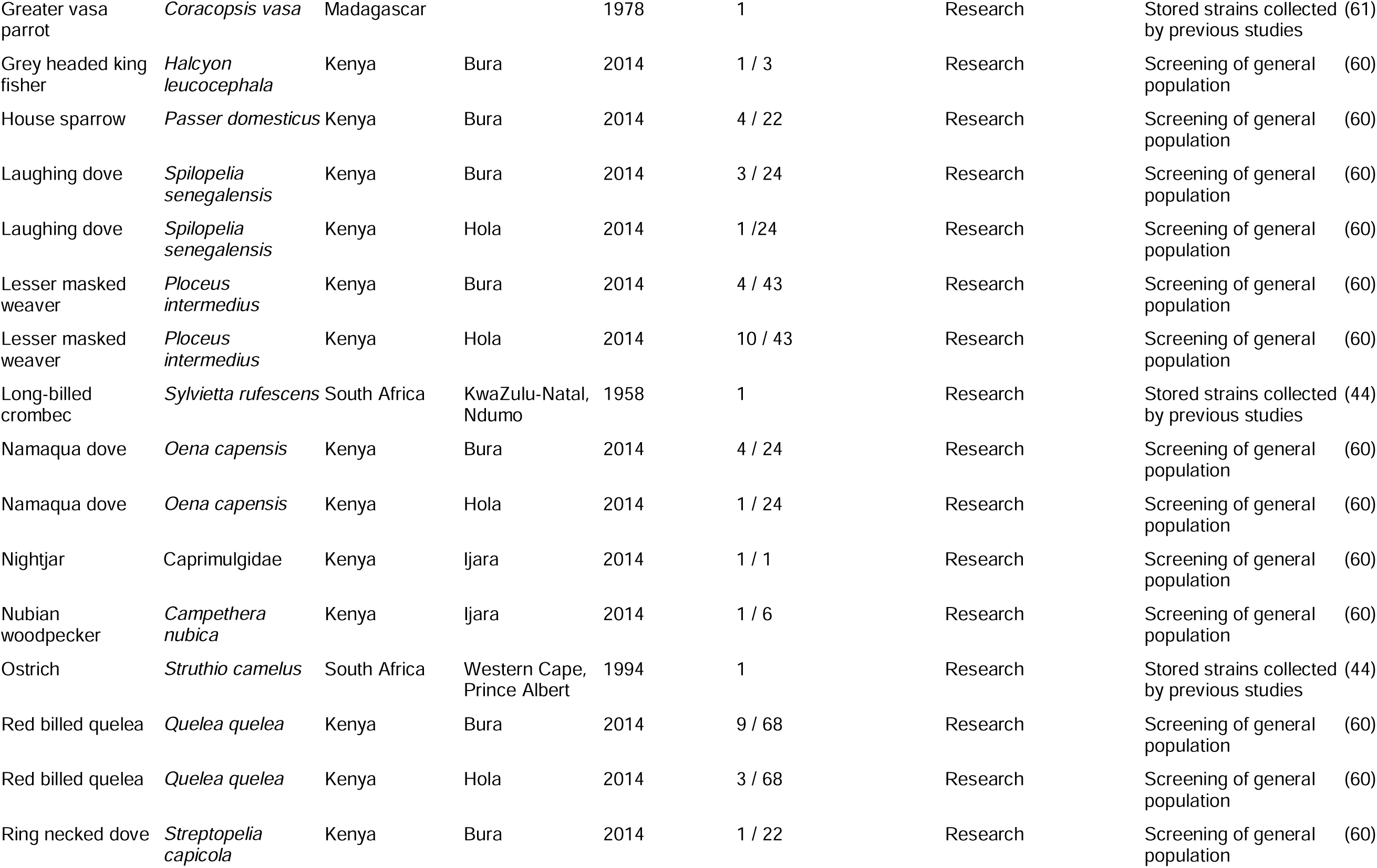
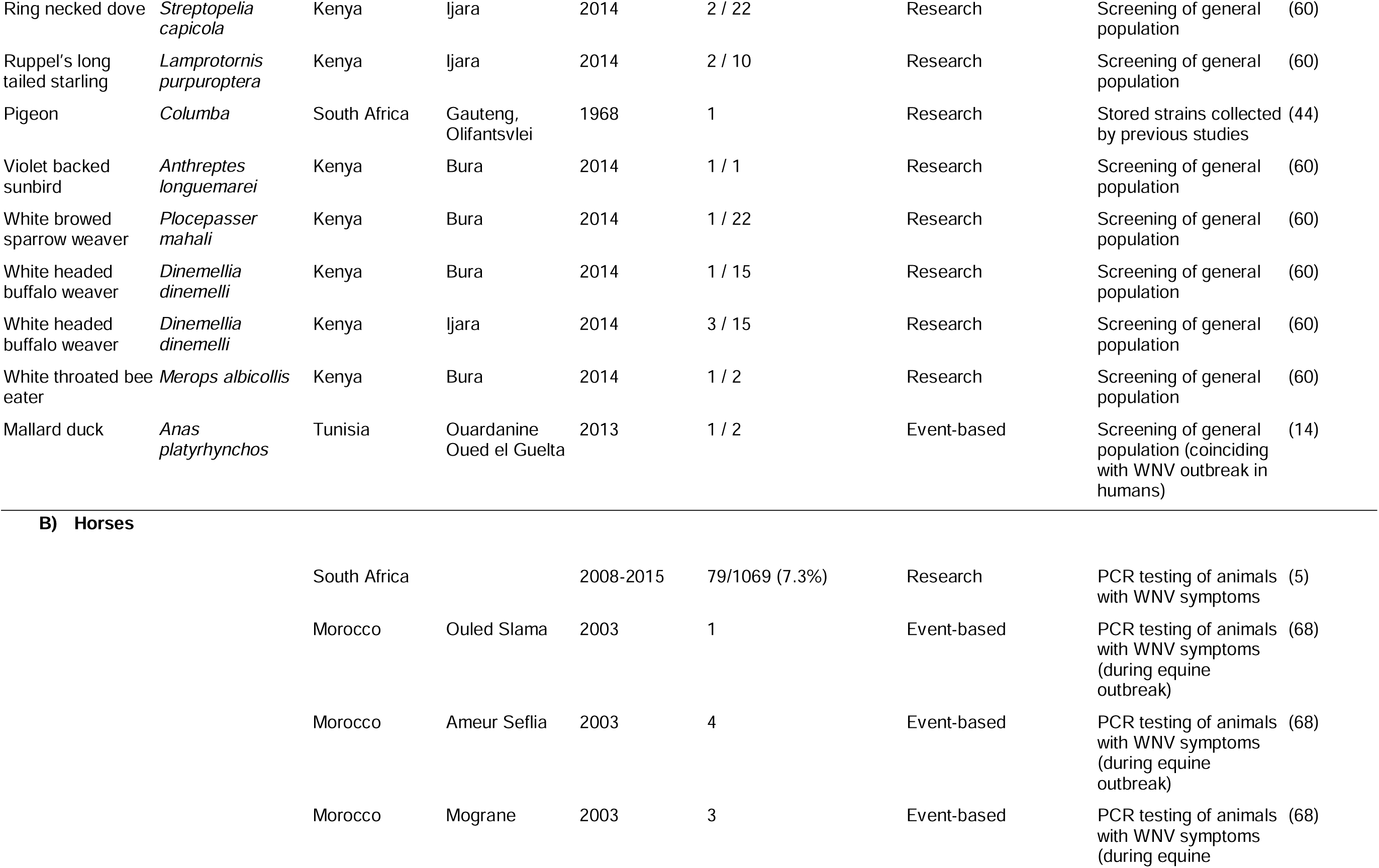
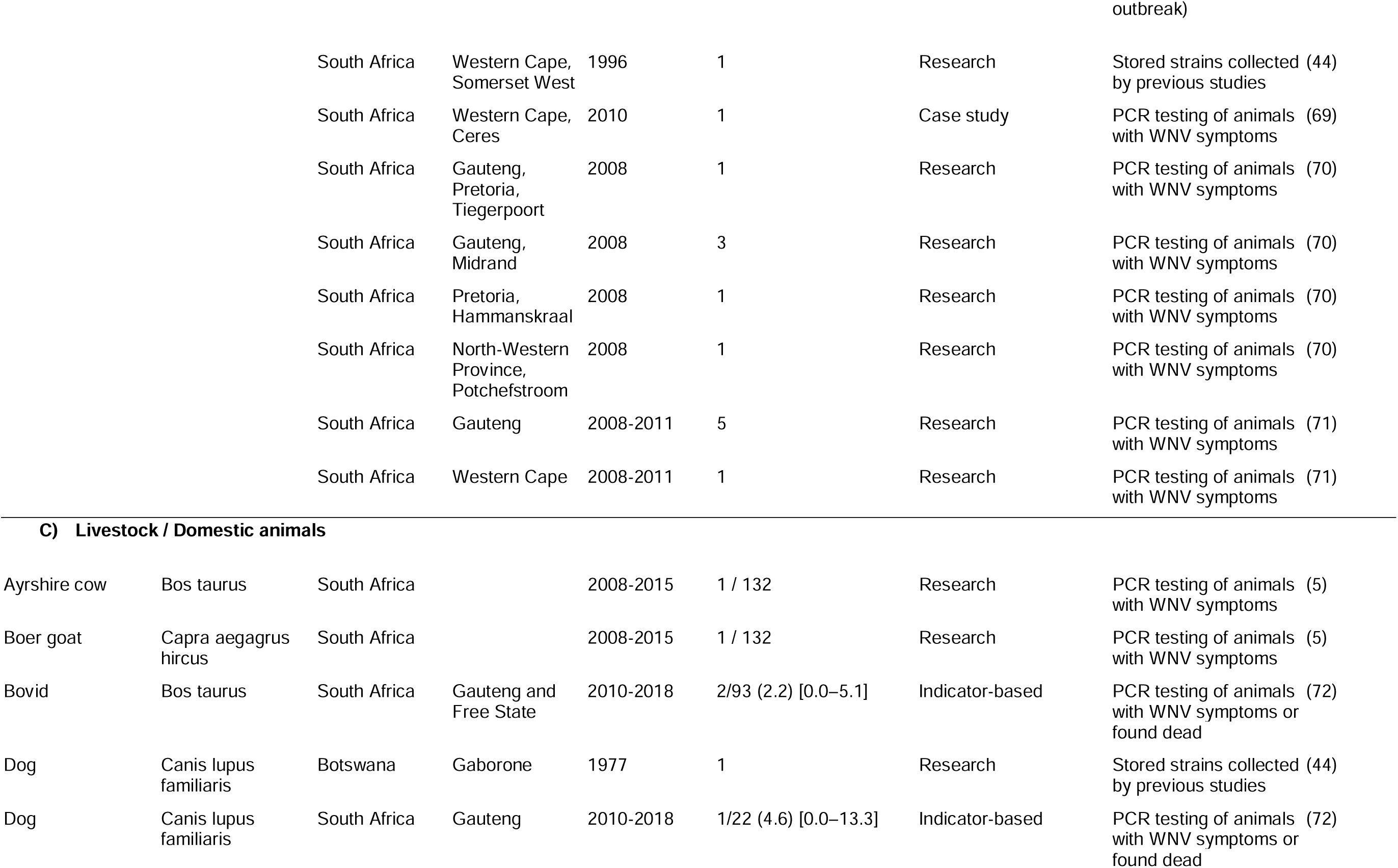
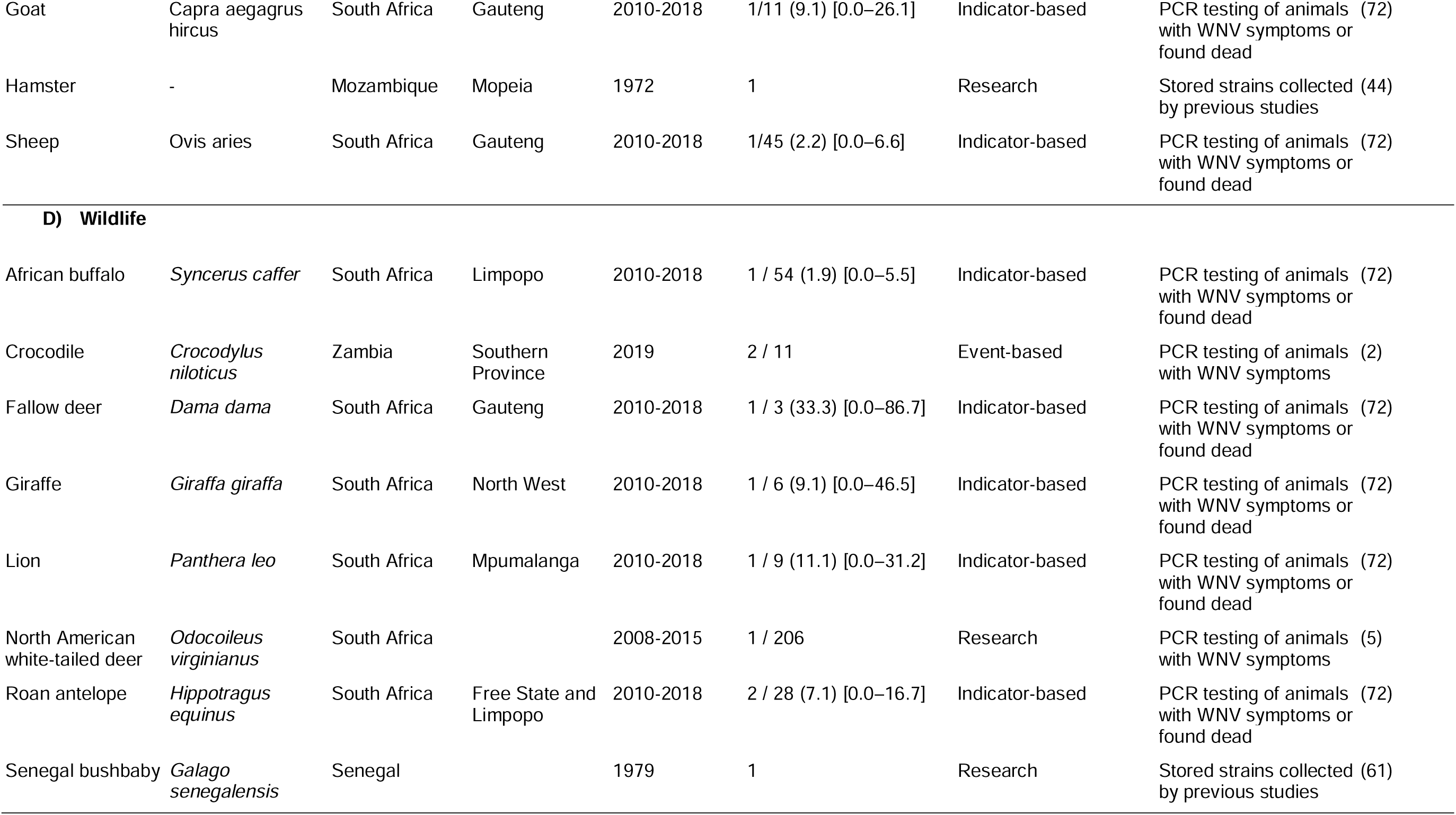
Positive molecular detections of West Nile Virus from wildlife and domestic animals in Africa. Sub-national location information is blank for studies from which sample collection location could not be retrieved. Positivity rate and 95% confidence intervals (CI) are provided if available from the primary study. Percentage positive is otherwise not provided as several studies did not utilise sampling methodology representative of populations and as such the calculated estimate would be biassed. Study type category and context of sample collection is provided for each study.

**Table 3:**
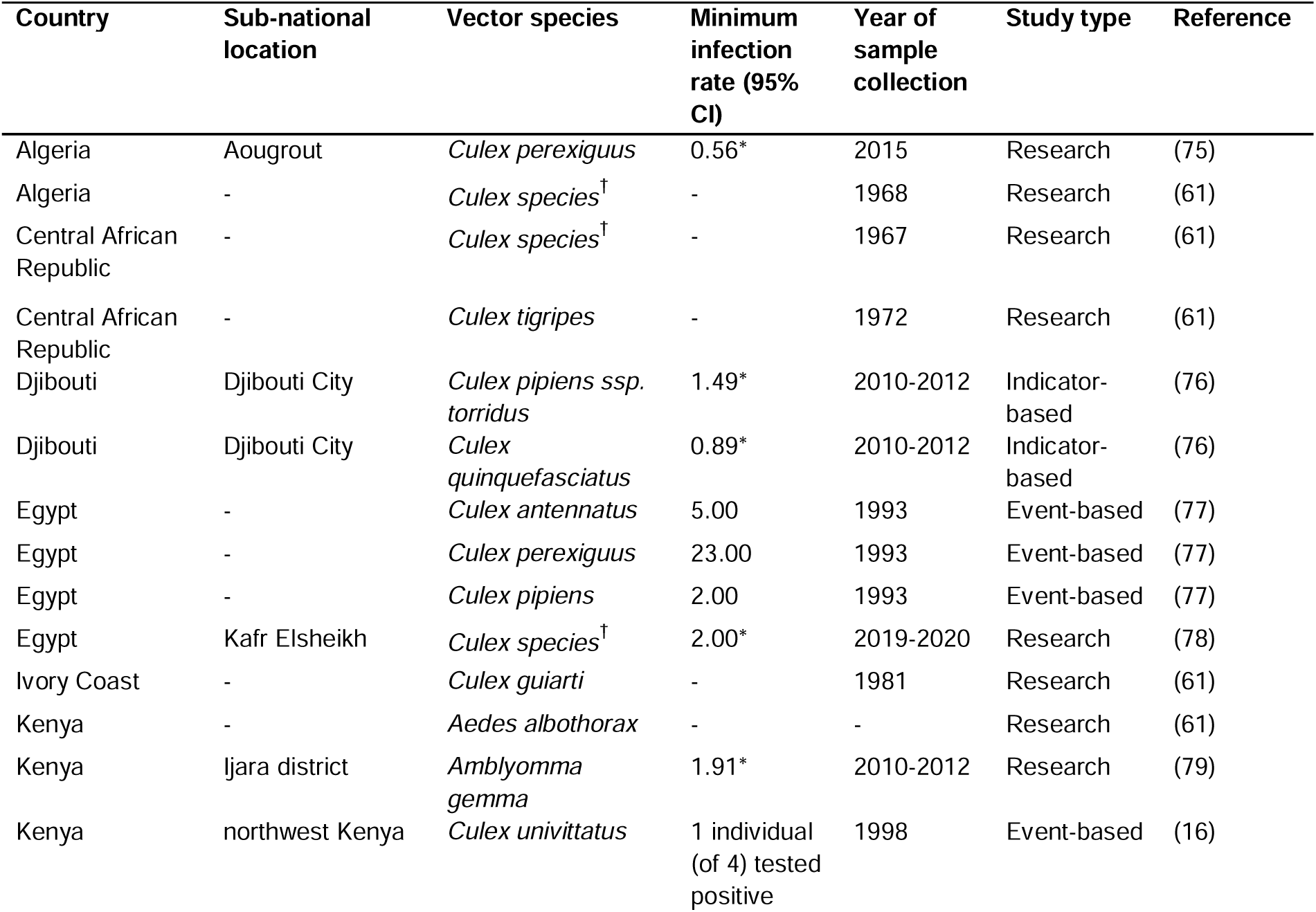
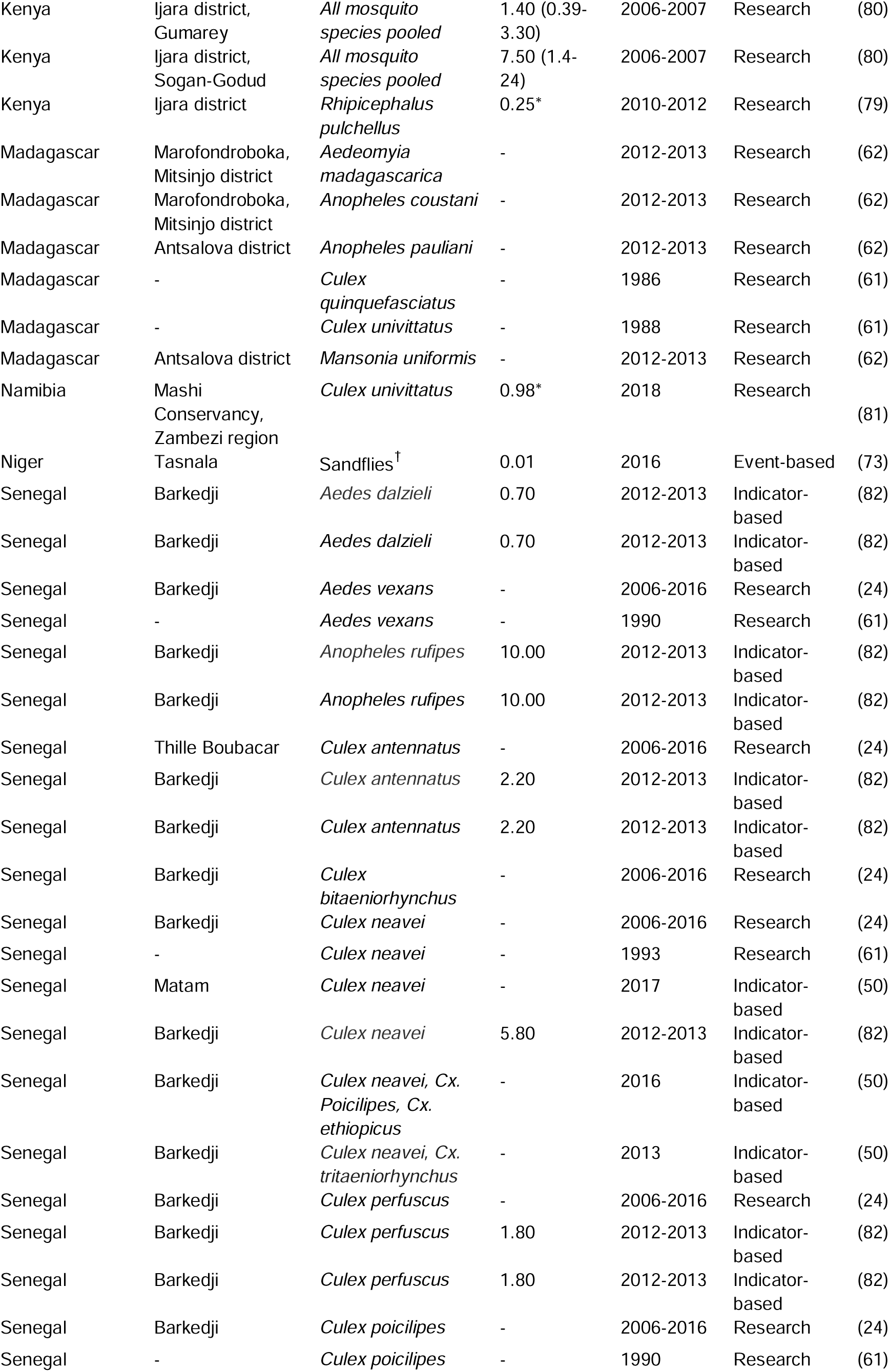
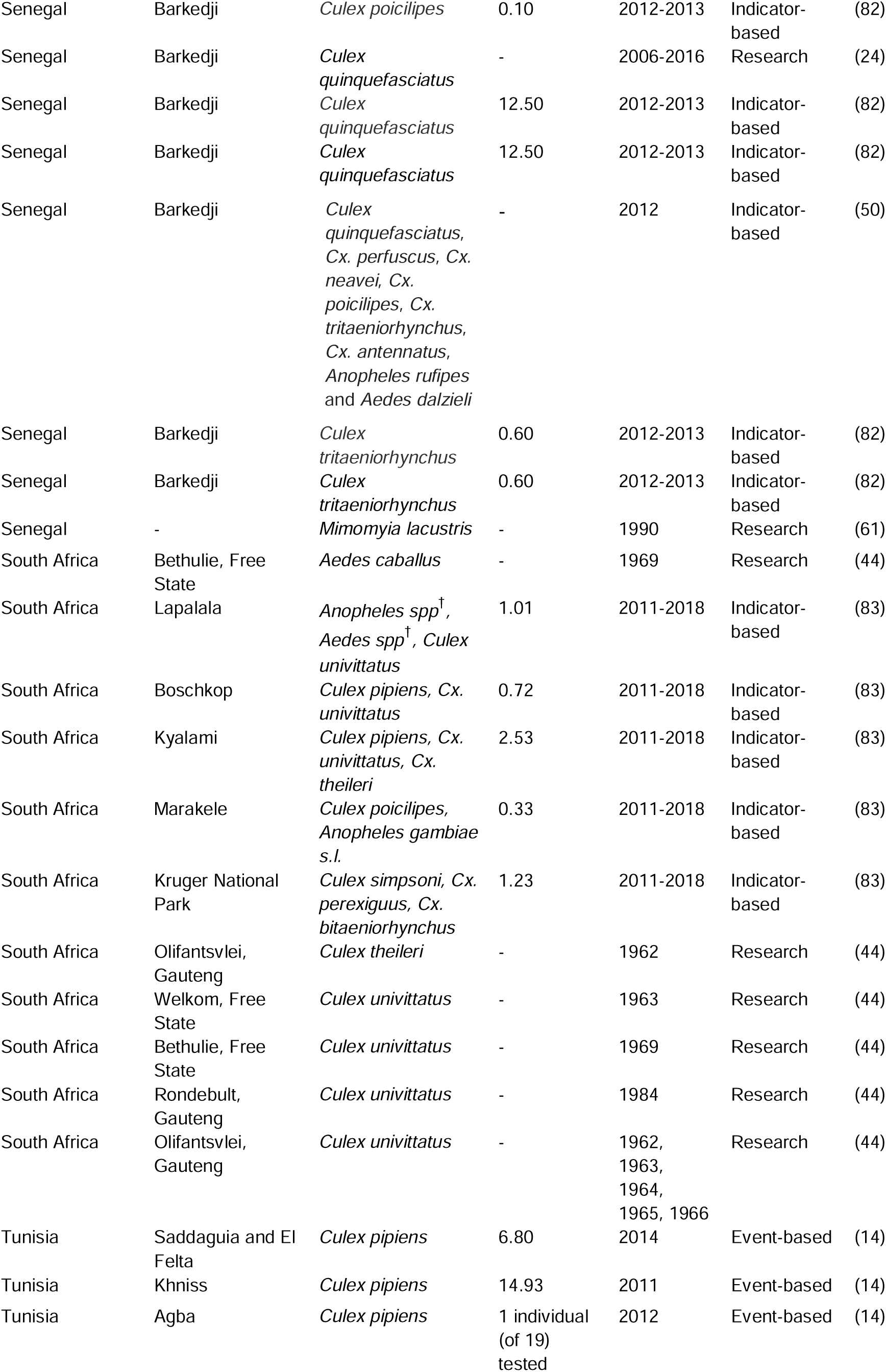
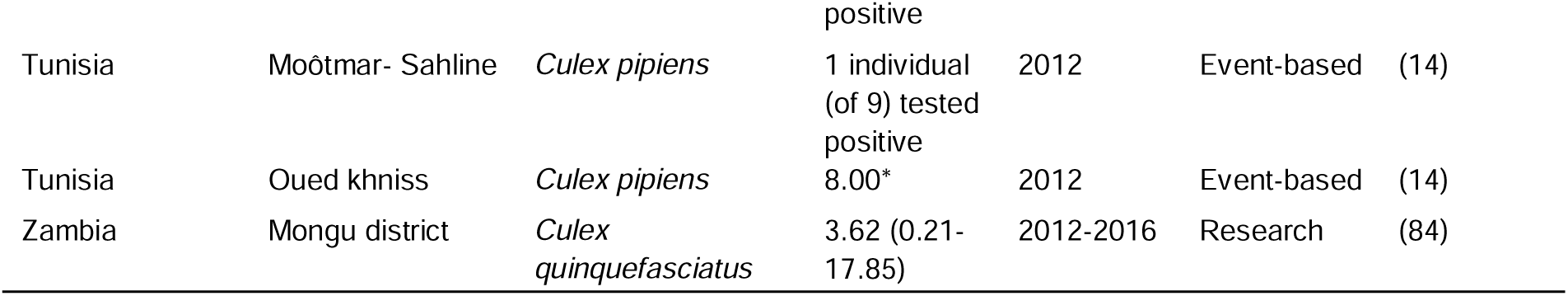
West Nile Virus isolations from mosquito and tick vector species per African country. Sub-national location of sample collection is provided or otherwise blank if the information could not be retrieved from the primary study. † denotes where taxonomic identification to species level was not provided. Minimum infection rate (MIR) and 95% confidence intervals (CI) are provided if available from the primary study. We calculated the MIR (74) if the primary study did not and the relevant information was provided. These values are denoted with an asterisk (*). Study type category is provided for each study.

Data on WNV PCR and/or sequencing in humans was available for only 12 African countries, of which eight reported positive detection of WNV (Table 1). Studies in Tunisia showed the highest percentage positivity for WNV and all infections were attributed to Lineage 1. This finding is unsurprising as studies included individuals with suspected WNV infection or confirmed WNV serology. Tunisia has experienced repeated outbreaks in 1997 (111 cases and 8 deaths) (40), in 2003 (112 cases and 9 deaths), 2012 (86 cases and 12 deaths), and 2018 (377 cases and 2 deaths) (41). PCR testing and sequencing was positive for WNV Lineage 2 in up to 9% of individuals hospitalised with acute fever of unknown origin in South Africa (42). A second study in South Africa detected WNV by PCR and sequencing in only 1 sample of individuals with suspected WNV infection, but found positive WNV serology in 19.4% of the cohort (43). Since these studies were conducted outside of recognized outbreak periods, the findings highlight WNV disease burden as underestimated in South Africa and an under-recognized cause of febrile and neurological disease in the country, particularly in children (42,43). PCR positive human infections were also found in Tanzania, Nigeria, Senegal and Sierra Leone when testing patients with fever of unknown aetiology and in one case study in Gabon, while WNV was not detected in human studies in Burkina Faso, Egypt, and Kenya (Table 1).

### Broad animal host range with low detection rates

Wild birds are reservoir hosts of WNV in endemic areas. Studies have detected WNV nucleic acids from 35 avian species in Africa (Table 2A). The majority of these species were sampled from areas in Tana River and Garissa in Kenya which is a known stopover for birds migrating to southern Africa from northern Europe (60). This is also a convergence point for domestic and wild animals for water providing opportunities for onward transmission of the virus (60). WNV has been isolated from a long-billed Crombec, sentinel pigeon, and an ostrich in South Africa from 1958, 1968, and 1994, respectively (44). Positive WNV detections were also found in a blackcap tchagra in CAR (61), a chicken (62), and a greater vasa parrot from Madagascar (61). All positive detections in bird species were outside of known WNV outbreak periods, except for one mallard duck sampled in Tunisia around the time of the 2012 outbreak in humans (Table 2A).

Horses are effective sentinel animals that are useful for estimating the risk of human infections in Africa (5). However, molecular studies in horses are available for only two African countries (Table 2B). We could find evidence of active surveillance programs for infectious pathogens in horses reported only in South Africa (5). An outbreak of Lineage 1 infections in horses was found in three locations in Morocco in 2003.

All detections of WNV in livestock and wildlife were from Southern Africa (from both Lineages 1 and 2) except for a Lineage 1 infection of a bushbaby from Senegal (Table 2C, D). We note a broad host tropism of WNV with detection in several species of livestock, domestic animals, as well as wildlife of the avian, mammalian, and reptilian classes. However, detection rates are generally low for animal hosts. Mammals are typically dead-end hosts as they fail to produce viraemia or are viraemic for a very short period (19), which greatly reduces the ability to detect the virus with PCR methods.

The majority of studies outlined in Table 2 tested samples from sickly or fatal animal cases while viral detections in bird hosts were found from studies screening the general population (presumedly non-symptomatic hosts). Reports of symptomatic birds in Africa are limited (63). However, in Europe and North America, where the disease was relatively recently introduced, several bird species present with neurological symptoms (64,65) with potentially devastating losses to local populations (66).

### WNV nucleic acids have been detected in 27 vector species in Africa

In Table 3, we present a synthesis of WNV detections from 13 African countries for six mosquito genera (*Aedeomyia*, *Aedes*, *Anophele*s, *Culex*, *Mansonia* and *Mimomyia*) and two tick vectors (*Amblyomma gemma, Rhipicephalus pulchellus*). The lowest infection rate of 0.01 was detected from sandfly vectors in Tasnala, Niger; it was found to be Koutango virus (Lineage 7) with a worryingly high virulence in mice (73). The greatest minimum infection rate (MIR - an estimate of the proportion of infected mosquitoes in a population) of 23.00 was from *Cx*. *perexiguus* in Egypts Nile Valley (Table 3). The virus was detected from its primary vectors: *Cx*. *pipiens* s.s., *Cx*. *quinquefasciatus* and *Cx*. *unitittatus*, in Djibouti, Egypt, Madagascar, Namibia, Senegal, South Africa, Tunisia, and Zambia.

### 127 administrative level one locations with WNV circulation that are lacking molecular surveillance

We collated the locations of WNV occurrences from reported cases, seroprevalence surveys, and other research studies for humans, animals and vectors. We used 72 literature sources to geolocate WNV detections in humans for 29 continental African countries, Comoros, Mauritius, Rodrigues, and Réunion Island. This data shows detection of the virus in humans in all regions of the continent, with high seroprevalence (>60%) from Algeria, Tunisia, Mali, Nigeria, Egypt, Sudan, DRC, Uganda, and Kenya (Figure 2A). Since 1937, we found a total of 301 human cases have been reported from 10 African countries and Réunion Island. WNV was detected from birds in 10 countries (Figure 2B). For all other animal species, we found evidence of WNV for 22 countries from 62 literature sources, of which 66% of the literature sources showed WNV detection in equine species (Figure 2B). We used 25 literature sources to gather locations of WNV detection from vectors for 13 countries (Figure 2B). Considering all host and vector species, WNV has been detected in 39 African countries (including Comoros, Seychelles, and Mauritius), the Canary Islands, and Réunion Island.

**Figure 2:**
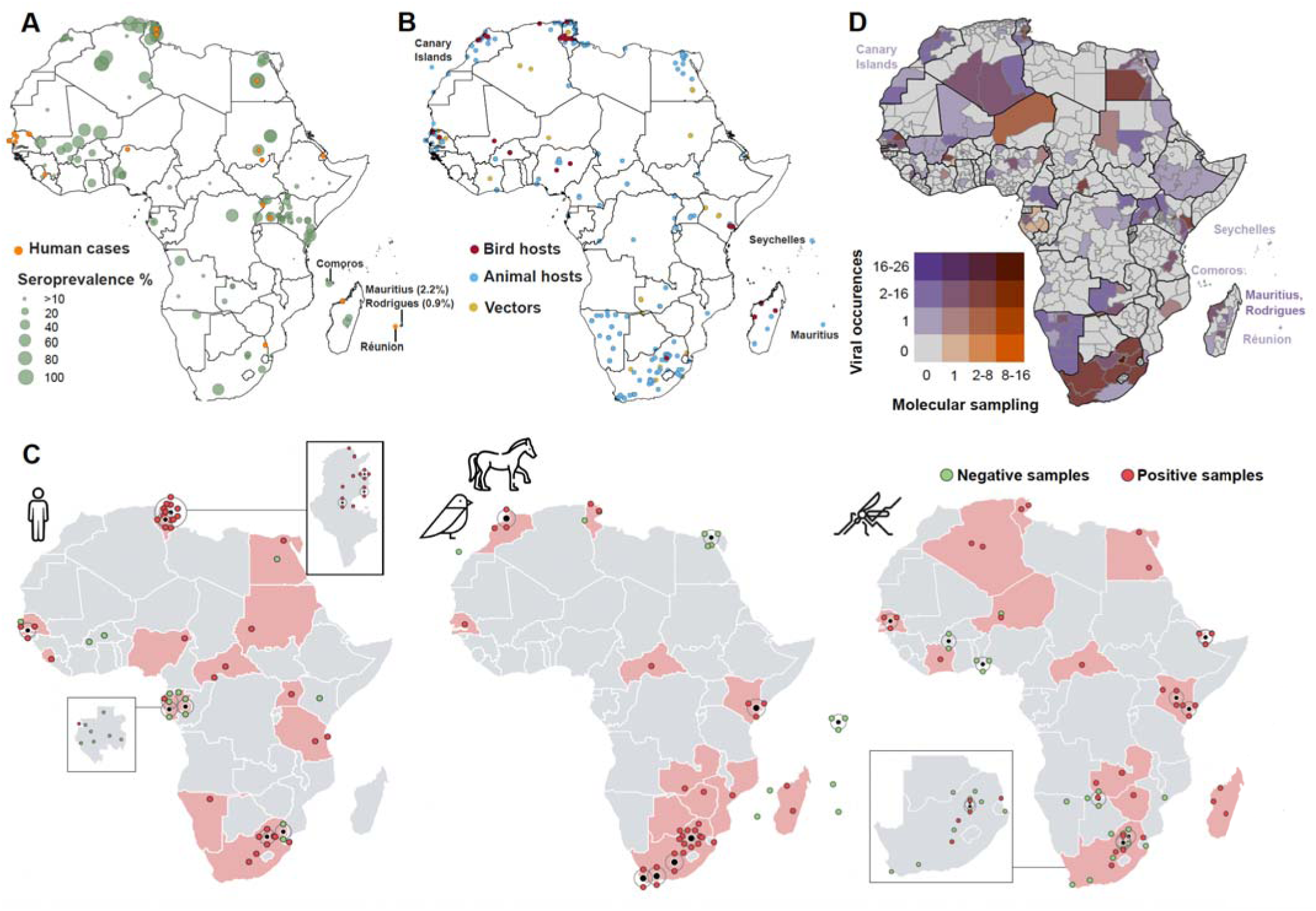
West Nile Virus circulation and molecular surveillance in Africa. A) Geolocations of West Nile Virus cases in human hosts (orange) and prevalence (% shown by circle size) from serological surveys (green); B) Georeferenced locations of viral detections (cases, deaths, seroprevalence) in reservoir bird hosts (red), dead-end animal hosts (blue), and mosquito and tick vectors (gold); C) Geographic distribution of positive and negative molecular detections (PCR and sequencing) of West Nile Virus from human hosts, animal hosts, and arthropod vectors. Light red shading indicates countries from which positive samples were collected. Point locations are shown with a point displacement method which indicates the locality of overlapping points with a black marker and the number of points at this location with coloured points on a black ring around the central marker. If detailed location data is not available, points are plotted in the centre of the country of collection; D) Bivariate map depicting WNV circulation (total number of detected viral occurrences from reported cases, deaths and seroprevalence surveys for all hosts) and the total number of molecular sampling locations on administration level one scale for all African countries. Locations were not mapped if they lacked detailed spatial information. Colour of Island names matches their designation of the bivariate colour scale.

During our review of molecular studies, we extracted fine-resolution sample collection locations to plot spatially explicit locations for studies that conducted PCR testing and sequencing of WNV (Supplementary information S6A). Detection of WNV by PCR has been reported by 13 countries and seven western Indian Ocean islands (Aride, Bird, Cousin, Europa, Juan de Nova, Réunion, and Tromelin Islands). The above-listed Indian Ocean Islands are mostly uninhabited by humans (except for Réunion Island). Gabon depicts the greatest dispersion of sample collection locations; mostly emanating from (47) which sampled non-malarial febrile cases from 14 areas across the country. Studies from 20 African countries and the Canary Islands collected samples for genomic sequencing (Supplementary information S6B). The study conducted in the Canary Islands tested wild cetaceans that suffered encephalitis or meningoencephalitis but did not detect WNV from 19 cases (85). Tunisia (n=8 studies) and South Africa (n=13 studies) showed the greatest extent of collection locations for sequencing studies. A large proportion of countries exhibit sparse sampling locations. We note the presence of WNV in Zimbabwe from a publicly available 225 nucleotide sequence (OL790153) lacking detailed location information, and an unpublished preprint from 2021 showing detections from crocodiles and mosquitoes (86).

In Figure 2C, we highlight detections of active WNV infections via PCR testing and/or genomic sequencing in 21 African countries, across all regions, for human and animal hosts and arthropod vectors. The vast majority of Tunisian studies tested human samples with positive detections in the north-western regions (9,45,46,56) with seemingly limited testing in the southern regions. All but one sampling location in Gabon returned negative results for human hosts; the positive detection was a lethal case of meningoencephalitis in a young man in Libreville (87). This is similar to Burkina Faso where human and vector samples tested negative (57) and *Aedes* mosquito populations screened for arboviruses in Benin returned negative results (88). We note records of reported human cases (Figure 2B), for which molecular work is absent, from Djibouti (89) and Madagascar (90). We display the temporal range of study publications captured by the review as well as the number of publicly available sequences per country (Supplementary information S7). There exists a comparatively high study and sequencing effort for Senegal, South Africa, and Tunisia. A limited number of sequences are publicly published for several countries, such as CAR, the Democratic Republic of the Congo (DRC), Egypt, Ethiopia, and Uganda. Several of these are the earliest sampled sequences from Africa which contribute invaluable information for phylogenetic investigations.

We link the circulation of WNV (total number of WNV occurrences) with the total number of molecular sampling locations, from all host and vector species, on a subnational administrative level (Figure 2D). This identifies a total of 127 administrative level one locations, and six islands and island archipelagos, with detected viral circulation but lacking molecular surveillance; particularly for many regions of Namibia, Ethiopia, Morocco, Western Sahara, and Mali. Regions with high WNV circulation and a relatively higher number of molecular sampling locations include seven provinces of South Africa (Gauteng being the highest), Garissa County of Kenya, and Monastir in Tunisia.

Considering the crucial role of birds in the transmission of this virus, we assume an increased risk of spillover to humans in areas in which birds spend an extended period of time (91). Key Biodiversity Areas (KBA) are areas harbouring significant biodiversity that are considered crucial for conservation of regional biodiversity (92). KBAs consist of Important Bird and Biodiversity Areas (IBAs) of international importance for birds (locations where birds breed or pause between migratory flights) identified by BirdLife International (92). These areas are also likely to harbour abundant vector populations. We identify regions with confirmed WNV detections that overlap with KBAs (map of African KBAs available in (92)) and areas of high human population density (https://www.kontur.io/portfolio/population-dataset/) that may be susceptible to large outbreaks. These areas include the Eastern Cape Province of South Africa, Western and North-Western provinces of Zambia, Afar and Oromia State of Ethiopia, four Governates of Egypt (Qena, Monufia, Kafr el-Sheikh, and Al Sharqi), Centre and East regions of Cameroon, and five regions of Morocco (Souss-Massa, Casablanca-Settat, Marrakech-Safi, Fez-Meknes, and Béni Mellal- Khénifra). In light of the great genetic diversity of WNV, widespread high human density, and KBAs littered across the region, all surveillance activities across the West African region would likely yield valuable information. Similarly, nationwide surveillance in Uganda is important, as the country from which it was first isolated so far has produced only three genomic sequences. Bordering Uganda is the Haut-Uele of the DRC, a further place of interest in which WNV circulates but which lacks genomic data. Research in the Great Rift Valley region in Ethiopia and Kenya may be of interest as there is a diverse number of bird species and potential for contact between humans and animals. Regions that may be of interest for zoonotic research due to high biodiversity and high WNV prevalence, but low human density, include Analamanga in Madagascar, North Kurdufan of Sudan, all regions of Namibia and Western Sahara, and Tassili N’Ajje biodiversity conservation area in Illizi District of Algeria (Figure 2D).

### Africa has produced the third-highest number of whole genomes in the world

We used publicly available genomic data to assess the temporal and spatial distributions of WNV lineages (Figure 3). Globally, North America has produced the greatest number of whole genome sequences (n = 3,012), followed by Europe (n = 701) (Figure 3A). Africa has the third-highest number of whole genomes (n = 63). In light of this, we assessed the laboratory infrastructure employed in the publications of the molecular review. We found that wet laboratory procedures were performed within the respective African country for the majority of studies (67.8%), with 28.8% employing an external laboratory, mainly located in the USA (35.3%) and Europe (35.3%) (Supplementary information S8). Regarding the molecular methods, 29.5% used solely RT-PCR methods, and 70.5% employed sequencing techniques, mainly of the non-structural protein 5 and envelope genes (Supplementary information S9). RT-PCR and Sanger sequencing have been consistently utilised in the studies, whereas next-generation sequencing methods were first used ten years ago with seemingly more regular use since 2021 (Supplementary information S10).

**Figure 3:**
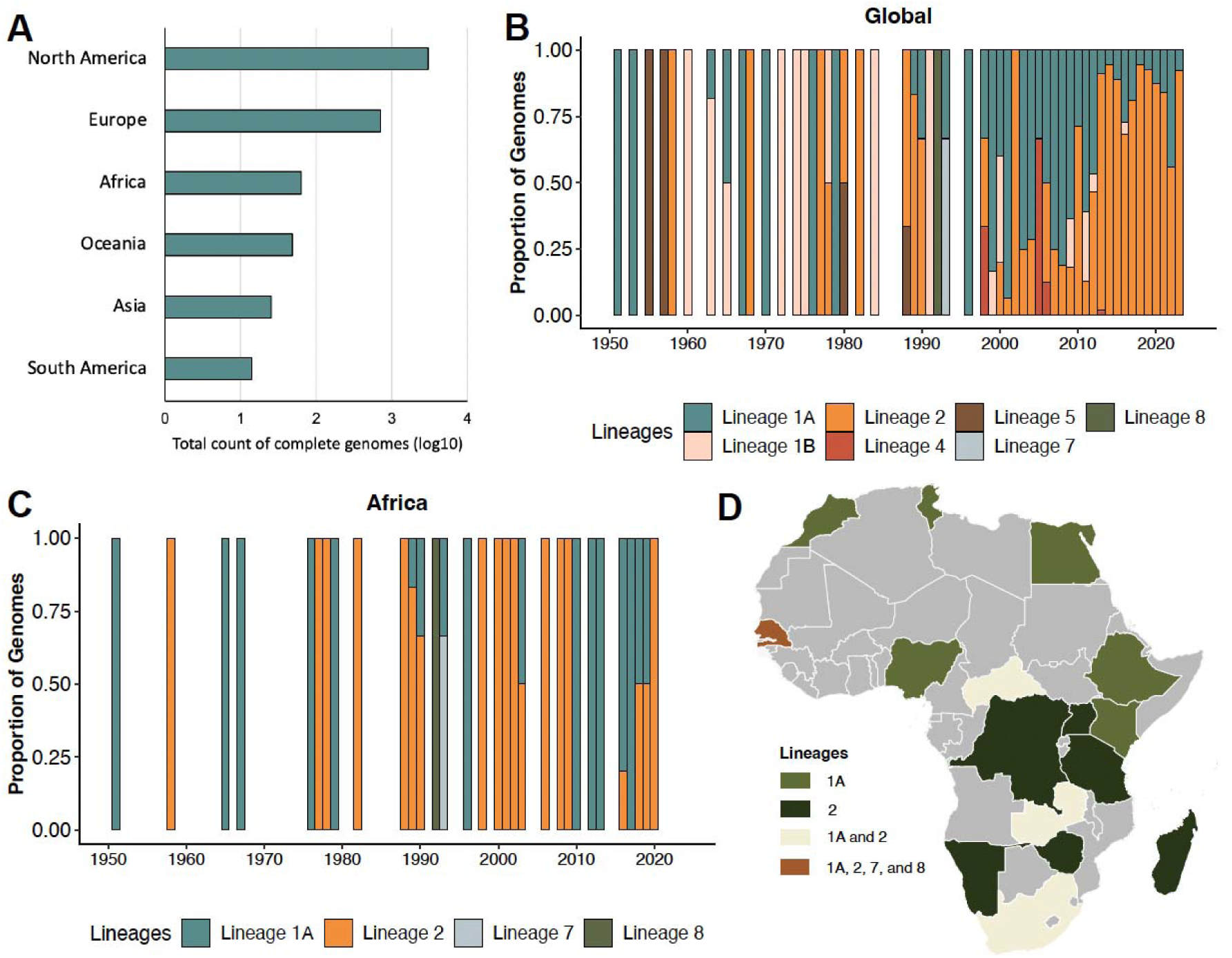
Global and African genetic diversity of West Nile Virus. A) Log transformed number of whole genomes per geographic region (retrieved from NCBI on 27 August 2024); B) Proportion of complete genomes per lineage of West Nile virus distributed over time from a global genomic dataset; C) Temporal distribution of proportions of complete genomes per lineage collected from Africa; D) Spatial occurrences of West Nile virus lineages in Africa from complete and partial genomes.

WNV encompasses up to nine genetically distinct lineages: lineage 1 (L1) to 9 (L9), although L7 has been classified as a distinct flavivirus, the Koutango virus (93). Globally, whole genomes are available for six lineages. Historically, L1A and B were the predominant circulating lineages but over the past two decades L2 has risen to dominance (Figure 3B). The change in lineage distribution has been attributed to the recent spread of L2 into the northern hemisphere and subsequent autochthonous transmission in Europe (94). There is high genetic diversity of WNV in Africa, with whole and partial genomes generated for L1A, L2, L8 and Koutango virus (Figure 3C). Similarly in Africa, L1A and L2 dominate the landscape of genetic diversity (Figure 3D). L7 and L8 have been sequenced only from Senegal demonstrating a wealth of genetic diversity in the country as well as successful genomic sequencing research and surveillance programs.

### Senegal is a dispersal hotspot for Lineage 1A

Our time-scaled phylogenetic reconstruction demonstrated a strong correlation from the molecular clock regression analysis for L1A (clock rate=0.00043, r^2^=0.94). Our spatially annotated phylogenetic analysis infers an almost global distribution for L1A with strains originating from Africa, Asia, Russia, the Americas, the Middle East, and Europe (Figure 4). The time from the most common ancestor (tMRCA) for all taxa in the L1A tree is estimated to be approximately 1915 (90% marginal probability distribution: 1472-1949), similar to the finding of (24). The basal clade of the tree (Cluster 1 - cluster nomenclature taken from (95)) consists of the oldest strains sampled from Egypt in 1951 (GenBank accessions AF260968 and EU081844) clustering with sequences from Europe, the Russian region, Israel, Tajikistan and Ethiopia collected in 1976 (AY603654) (Figure 4A). The tree branches into larger clades of Asian and Russian isolates (Cluster 3), North and South American strains, and European sequences (Cluster 4); all of which are rooted by older African taxa from Senegal (GQ851606 collected 1979) and Nigeria (GQ851607 from 1967) in Cluster 5. The crown of the phylogeny depicts transmission within Europe (with the earliest clade divergence dated 1984, 90% CI: 1843-1996). African strains from Senegal (2012-2018) and Morocco (AY701412-3; 1996 and 2003) as well as an Israeli sequence (2000) are basal within this cluster. Outside of this cluster are several diverse sequences from Africa, Europe, Middle East and Russian regions.

**Figure 4:**
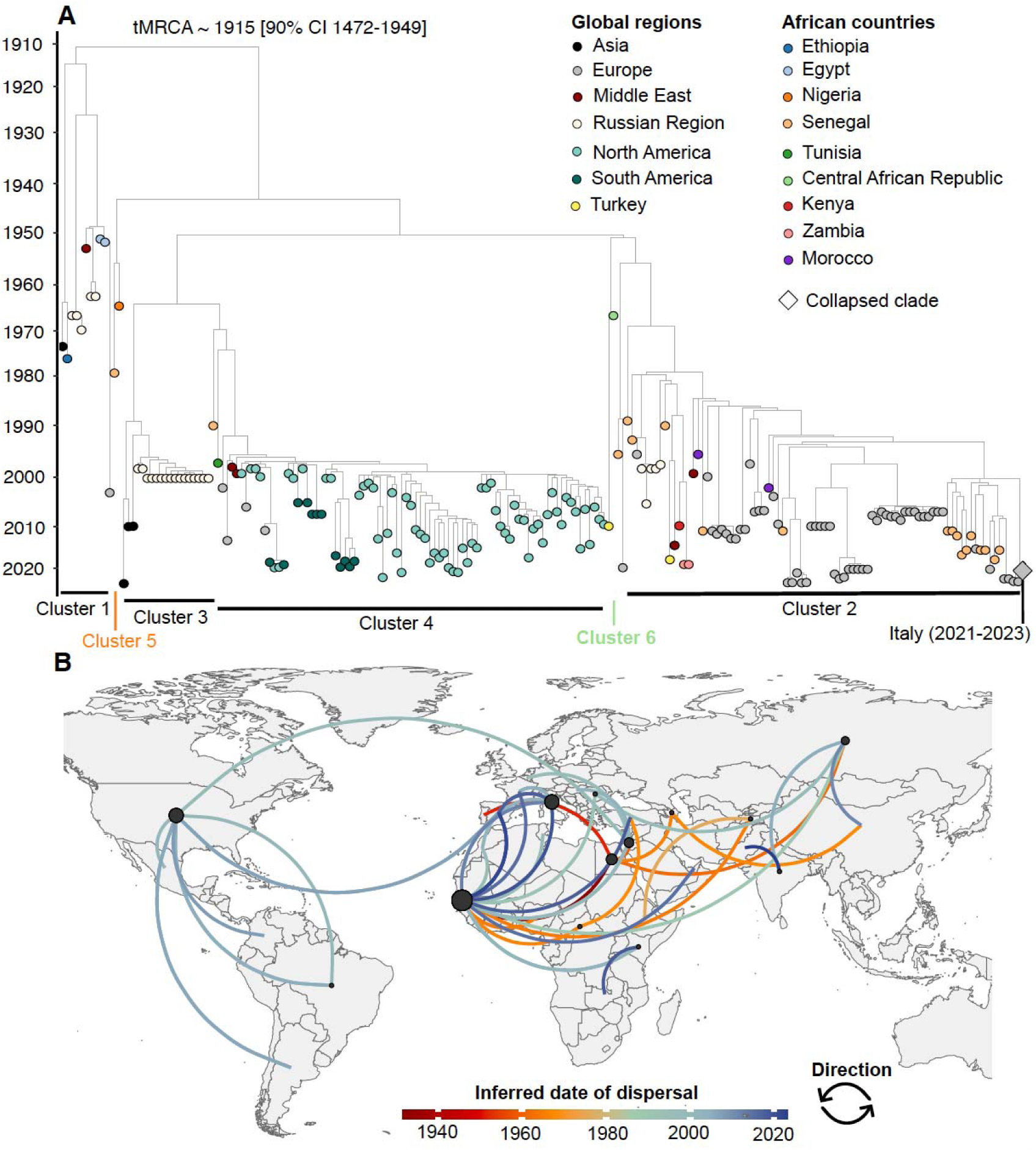
Lineage 1A reconstruction. A) Maximum likelihood time-scaled tree depicting global diversity of West Nile Virus Lineage 1A strains. Tree tip colours represent global regions or African countries from which strains were collected; B) Map illustrating the spatiotemporal dissemination patterns of West Nile virus L1A across and within Africa, Europe, Asia, and the Americas. Inferred dispersal pathways are coloured by the mean date of viral transitions for each route; earlier transitions are shown in red to later transitions in dark blue. Circles show the inferred number of exports per country (largest circle represents 20 exports; smallest circle depicts one export). The direction of movement is shown from the origin (black dot) to the destination with curved lines anti-clockwise in the direction of the curve.

Our global phylogeographic reconstruction of L1A inferred a West African origin as the earliest inferred transition was between Senegal and Egypt in 1931 (Figure 4B). We see several early dispersal events from Africa to other global regions: from Egypt to Portugal (in 1951), Israel (1959), Russia (1962) and Azerbaijan (1965); from Senegal to Tajikistan (1964 - with transmission back into Africa via Ethiopia in 1976), Italy (1985), Russia (1989), Israel (1991) and Romania (1992). Within Africa, L1A strains are estimated to have dispersed from Senegal to Nigeria in 1965, Central African Republic in 1966, Morocco (1996), Tunisia (1997), Kenya in 1998 and 2001 (Figure 4B). The most contemporaneous transition within Africa was from Kenya to Zambia in 2018.

Overall, we found at least four dispersal events out of Africa to both Europe and Asia, with three transitions back into Africa from Europe and one from Asia (Figure 4B). We infer three introductions from continental Africa into Middle Eastern countries with seven transitions within Africa. The greatest number of African exports originated from Senegal followed by Egypt, whilst most European exports arise from Italy and the USA for exports in the Americas.

### The first intercontinental dispersal of L2 strains occurred from Africa to Asia

Our time-scaled phylogenetic reconstruction showed a modest clock correlation for L2 (clock rate=0.00026, r^2^=0.53). The estimated tMRCA for all taxa in the tree is ∼1689 (90% CI: 1598-1899), which is earlier than the estimate of (24) that falls within our confidence intervals. One of the two oldest sequences, collected from South Africa in 1958 (HM147822), forms the most basal clade 2b and groups with a strain collected from Namibia in 2020 (Figure 5A). Two Madagascan strains, collected in 1988, form the clade 2c that diverged ∼1940 (90% CI: 1644-1976). Genetically divergent strains from Madagascar (1978) that form clade 2a were dropped from the phylogeny as they did not follow the molecular clock. The other 1958 South African strain (EF429200) clusters in clade 2d with strains from South Africa and CAR (divergence ∼1972, CI: 1815–1987), rooted by a DRC sequence (HM147824). Russian strains, along with sequences from Romania, Italy, and Hungary, diverge from basal African sequences ∼1990 (90% CI: 1893-2002). Senegalese sequences (1989-2006) group with a 1980 Ukraine strain at the base of the Western and Central European transmission clade (divergence dated 1971, 90% CI: 1835-1980). The tree extends into a large cluster from 16 European countries, collected from 2004 to 2023, and is rooted by the Hungarian sequence MZ605398. (Figure 5A).

**Figure 5:**
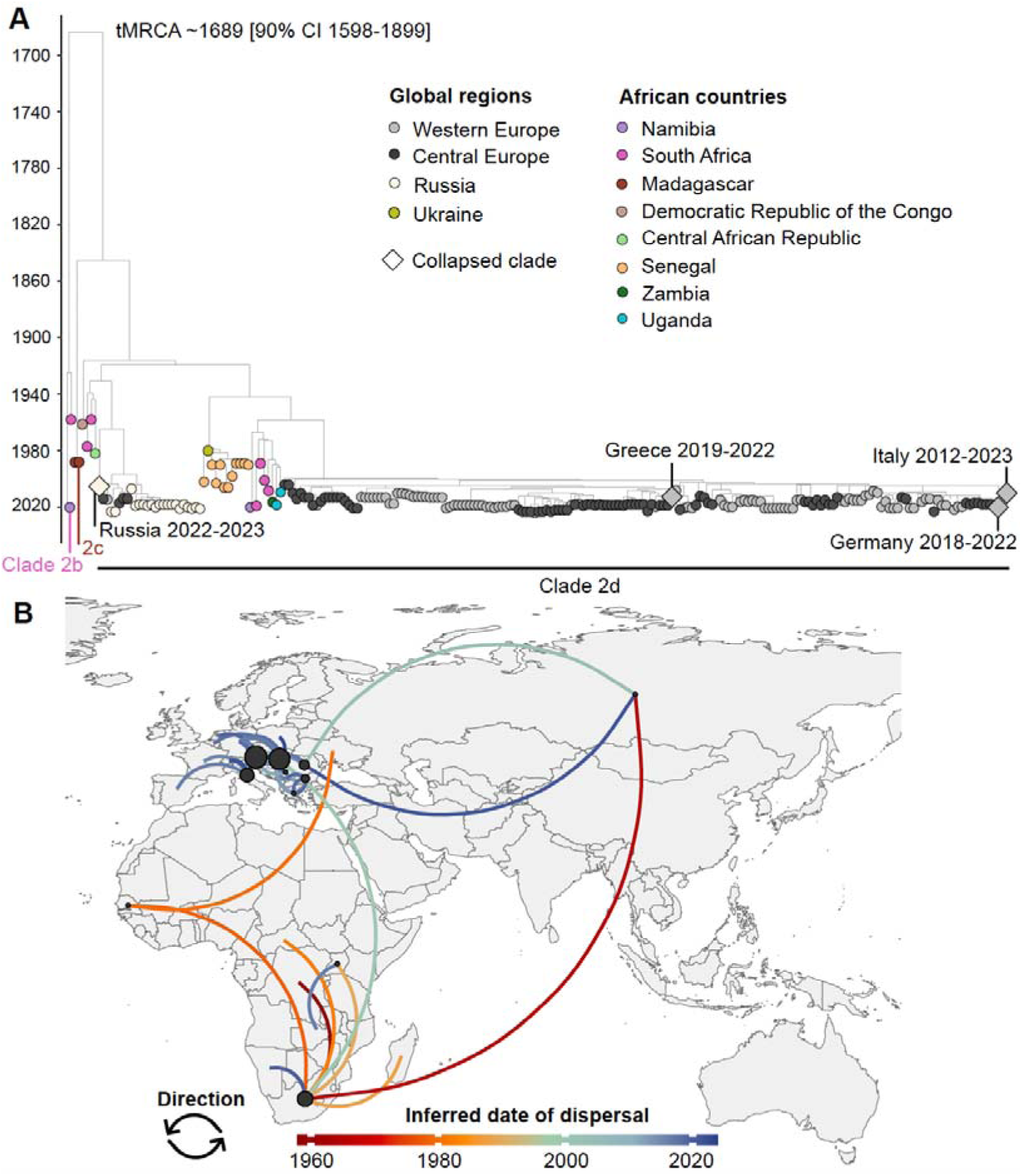
Lineage 2 reconstruction. A) Maximum likelihood time-scaled phylogeny showing the relationships between West Nile Virus Lineage 2 genomes. Colours of tree tips represent the European regions or African countries of sample collection. Diamonds display large clades of single origin sequences (collapsed for visualisation purposes); B) Geographic diffusion pattern of Lineage 2 genomes between and within Africa, Asia, and Europe. Inferred dispersal pathways are coloured by the mean date of viral transitions for each route; earlier transitions are shown in red to later transitions in dark blue. Circles show the inferred number of exports per country (the largest circle represents 23 exports; smallest circle depicts one export). The direction of movement is shown from the origin (black dot) to the destination with curved lines anti-clockwise in the direction of the curve.

The phylogeographic reconstruction shows South Africa as the primary source of L2 transitions, with the highest estimated exports (Figure 5B). Early transitions within Africa are inferred from South Africa to DRC (1958), Senegal (1978), CAR (1982), Madagascar (1987) and Uganda (1989). The first intercontinental dispersal, from Africa to Asia, occurred in 1964, with no evidence of a return to Africa. Early estimated transitions to Europe were from Senegal to Ukraine in 1980 and South Africa to Hungary (1999). Austria and Hungary demonstrate the greatest number of inferred exports, all of which were to neighbouring European countries. There is one exchange between Asia and Europe with an initial transition from Russia to Romania. Intra-African transitions are more numerous than intercontinental exports from Africa, with more recent dispersal occurring from Uganda to Zambia (2016) and from South Africa to Namibia (2020).

## Discussion

Although WNV is included on the WHO list of priority pathogens with a high risk of causing public health emergencies of international concern (7), we demonstrate the significant knowledge gaps on the true burden of disease, molecular epidemiology, and distribution of WNV in Africa. Our analysis revealed evidence of WNV circulation for 39 of 55 African countries. For all remaining countries, no studies or reports were found that had investigated WNV circulation, except for Burundi wherein one study assessed arboviral infections in humans and did not detect WNV antibodies (96). It is important to note that there is significant serological cross-reactivity among flaviviruses (97) and multiple antigenically related flaviviruses co-circulate in several African regions. When assessing the geographical extent of WNV circulation in Africa, we collected data from studies reporting WNV detection from a range of serological methods. Considering the problem of cross-reactivity, a limitation of our study is that WNV circulation may be overestimated. Molecular data is available for 24 countries, but most lack data for all hosts and vectors: 12 countries reported on humans, 11 reported on animals, and 13 on vectors. Senegal, South Africa and Tunisia are the only countries with molecular data for humans as well as vectors and animal hosts. At the time of writing, only 63 genomic sequences originating from 16 African countries are publicly available.

It is essential to improve our understanding of the WNV disease burden and distribution in Africa to direct the development of diagnostic, treatment and prevention strategies. Considering the increasing global spread of WNV, this is relevant not only for the African region but globally. We show that Africa has been the birthplace of internationally important lineages and is globally interconnected due to long-distance bird migrations. While much attention has been paid to the recent introduction and spread of WNV across North America and Europe, comparatively limited work has been done in Africa, despite epidemics occurring as early as 1939 in Uganda, Sudan, DRC, and Kenya (98,99). We have identified three key challenges, which are in line with the WHO technical brief on research prioritization for pandemic and epidemic intelligence (100), that we believe will be useful starting points to address this knowledge gap on WNV in Africa.

### Need for low-cost diagnostics at the point of care

We draw attention to the pressing need for low-cost diagnostic testing for WNV that can be performed at the point of care. WNV often presents with non-specific febrile symptoms, making clinical recognition of the infection challenging. Currently, diagnosis of arboviral infections in Africa primarily relies on serological assays, which are often directed toward other more commonly suspected endemic arboviruses, such as dengue and yellow fever. Interpretation of arboviral serology is becoming increasingly difficult due to the global expansion of arboviruses, which induce antibodies that may cross-react in serological assays (101). Confirmation of positive serology through PCR testing is recommended; but is typically limited to national reference laboratories or public health institutes, resulting in long turnaround times. Also, periods of viremia are very short so patients often present to clinics too late for detection of viral RNA. In resource-limited settings, diagnostic testing for WNV– which is often self-limiting and currently lacks antiviral treatment options–is not prioritized. However, diagnostic testing for viral infections, including WNV, could direct therapeutic management. For example, it could facilitate early cessation of empiric antibiotic treatments, as seen for viral respiratory PCR panels (102).

The limited diagnostic capacity for WNV likely contributes to an underestimation of its human disease burden (103). For example, although South Africa has the highest number of publications containing WNV molecular data, the available data derives mostly from research studies. One such study demonstrated WNV is an underrecognized cause of neurological and febrile illness in South Africa (42). Despite it being a notifiable condition in South Africa, WNV cases are considered largely under-reported with approximately 5-15 cases reported per year via passive submission of suspected arboviral cases for diagnoses (19). Such under-reporting is also a result of a lack of clinical awareness of WNV as a cause of neurological disease and its associated disease profiles. WNV and other arboviruses should be considered as a cause of neurological disease with syndromic testing.

Access to affordable point-of-care diagnostics is essential for rapid early detection, increased clinical awareness, and addressing knowledge gaps regarding the disease burden in Africa. A review of the latest advances in arboviral diagnostics highlights several isothermal nucleic acid amplification technologies with potential as point-of-care diagnostic tools (104). However, for such devices to be adopted in resource-constrained settings they must be capable of accurately diagnosing several different arboviruses. This capability would support the detection of arboviruses that are less routinely tested for. Improved diagnostic capacity for WNV will also facilitate the development of integrated genomic surveillance systems, a second key challenge.

### Need for an integrated surveillance approach

We show evidence of WNV detection from a broad host range, multiple vector species, and many geographical regions in Africa. WNV has been isolated from at least one host species in all African regions indicating how adaptive the virus is to a wide variety of ecosystems (91) with potential for spillover to humans. We highlight the need for an integrated One Health surveillance approach for WNV. Such a surveillance system may fill in knowledge gaps of WNV disease burden for all host species and guide the design of effective public and animal health interventions. Such an integrated One Health surveillance approach in a WNV endemic area has proven economic returns with reduced public health costs (105). However, we recognise that establishing a comprehensive surveillance program (encompassing public health surveillance, equine surveillance, xenosurveillance and testing of many animal groups) is challenging and very costly, and may not be immediately feasible for many African countries. In such cases, prioritizing equine surveillance programs may be a strategic allocation of resources as such programs have demonstrated success as an early warning system for human outbreaks (106).

Here, we show diverse disease and genomic surveillance capacities across the African continent. When evaluating the molecular data from the continent, three countries stand out as the leading WNV genome generators: Senegal, South Africa and Tunisia. Successful sequencing efforts from Senegal are likely due to the country’s advanced arboviral disease surveillance capacity, well-functioning case notification and case management systems, and high preparedness for outbreaks (107). South Africa demonstrates well-functioning disease surveillance and case notification systems for arboviruses but may be hampered by the lack of a national programme on arboviral disease surveillance, and national surveillance programs of animals and vectors (107). It seems that South Africa’s large number of WNV sequences has been driven by academic research groups and the National Institute for Communicable Diseases of South Africa (5,42,72,83). The volume of published sequences from Tunisia likely emanates from their integrated One Health surveillance approach for WNV(108) which includes monitoring of human meningitis cases, clinical equine encephalitis cases, and passive surveillance in birds (109). A potential reason for limited surveillance efforts from other regions is that WNV is a notifiable medical condition in only a few African countries: Egypt, Morocco (110), Tunisia (109), Algeria (111), South Africa (https://www.nicd.ac.za/nmc-overview/), Sudan, South Sudan, Sierra Leone, Cameroon and Algeria. In the remaining regions where it is not a notifiable condition, WNV is likely not tested for due to national prioritisation of other epidemic-prone diseases (112).

There are several barriers to integrated disease and genomic surveillance of arboviruses in the African context. For example, viral detection in vectors is challenging and costly as it requires sampling large numbers of mosquitoes, such that a lack of viral detection may be due to under sampling rather than a true absence of the virus (113). The lack of pathogen detections, possibly due to under sampling, may make it difficult to motivate for further funding and sampling. Such challenges are further exacerbated by limited medical entomology capacity outside of malaria-prone areas, insufficient training, resources, mentorship, sharing of data, and collaboration (114). As became evident during the SARS-CoV-2 pandemic, many African countries lacked adequate sequencing infrastructure and local capacity for genomic surveillance (115). However, the SARS-CoV-2 pandemic resulted in a continent-wide increase in genomic sequencing infrastructure (115), which should now be leveraged to increase the available genomic data for many other infectious diseases.

Indeed, several resources in the region could be leveraged to support arbovirus surveillance such as the established capacity for malaria testing and control (107). Significant progress has been made by the establishment of networks that support arbovirus research and surveillance (114). Additionally, to address shortfalls in technical abilities and support, there exist several capacity-building and training initiatives for the African continent (116–118). These collective efforts underscore the potential for enhanced arbovirus surveillance systems and highlight the significant strides already made on the continent.

### Need to integrate diverse datasets for an enhanced understanding of WNV emergence

Data generated by disease and genomic surveillance systems are not only important for an informed public health response to an outbreak, but are also crucial for analytical techniques and modelling methodologies to predict potential outbreaks for preventive public health measures. We underscore the need to generate, share, and integrate diverse datasets to improve our understanding of WNV emergence and re-emergence. Below, we provide recommendations for the generation of novel and diverse datasets for impactful contributions to our understanding of WNV.

To further our understanding of WNV transmission, dispersal, and evolution via genomic analytical techniques, we recognise the need for genomic data from more geographical locations in Africa. However, in resource-constrained settings, it is vital to identify regions for targeted surveillance to inform resource allocation in the most impactful way. Here, we identified regions with confirmed WNV circulation but lacking molecular surveillance. We also note the lack of molecular sampling and sequencing for Lesotho, Eswatini, and The Gambia; countries nestled within the borders of the highest WNV genome-producing countries, as well as for those neighbouring them. There also appears to be a general absence of case reporting and seroprevalence surveys from these countries. However, risk mapping shows high potential for WNV circulation (91), with significant cross-border travel between endemic countries and their neighbours (119,120), such that we suspect transmission is occurring but has not yet been sampled or detected. Disease surveillance and genomic data generation from these regions is likely to be impactful for our understanding of WNV transmission in Africa. Additionally, the generation of more genomic data is essential to advance vaccine development, which remains lacking for preventing human WNV infections.

Even though publicly available African WNV genomic sequences are limited, our phylogenetic analyses allow us to understand that Africa is the main source of diverse viral genotypes (95). Our phylogeny of L1A depicts six previously delineated clusters (95), with all but cluster 3 containing basal African isolates. The same is true for all clades of L2. This is suggestive of recurring spread of different genotypes out of Africa and indicates a source-sink dynamic with Africa as the source. Similarly, our phylogeographic results allow us to identify several transitions between other continents and back transitions into Africa that indicates continual viral movements between global regions (121). The interconnectedness of most global regions in WNV transmission is attributed to the annual long-range migrations of wild bird hosts (122–124). Considering the general paucity of WNV genomic data and the continually sustained bird migrations transporting the virus, it seems likely that only a minority of these viral exchanges have been detected. Despite the importance of migratory birds in the dispersal dynamics of this virus, primarily risk modelling has been done to assess viral spread via African flyways (122–124). We show molecular testing for WNV in bird hosts is limited to five countries and recommend further work be done to better understand dispersal of WNV via bird amplifying hosts to and from migration hotspots in Africa.

Phylogeographic analysis showed the earliest inferred export and the greatest number of exports of L1A was from Senegal. This raises the question of why this country is a dispersal hotspot for WNV. We postulate that it’s due to persistent viral circulation paired with the geographic and ecological roles of Senegal in bird migration. Sustained endemic circulation is facilitated by high prevalence of several bird species (13) and the abundance of competent mosquito vectors (50,82,83). Senegal and Europe are well-connected via the largest land bird migration network, the Afro-Palaearctic Bird Migration System (125). The majority of migratory birds depart the Sahel region for Europe in March–April (126), corresponding to the warm and dry season in Senegal. As water sources dry up, birds congregate in areas with the last remaining water sources. The highest WNV vector abundances were found in barren and temporary ponds in Barkedji, Senegal (127). Such congregations of both bird hosts and mosquito vectors may create conditions of high transmission risk immediately prior to the departure of birds for their return to Europe. However, viraemia is transient in birds (5-7 days) (128) and migrations can take from two to several weeks (126). We postulate that infected ticks carried by trans-Saharan migrant birds from epizootic areas may facilitate long-distance introductions (129) as transstadial transfer of the virus has been demonstrated with onward transmission for an argasid tick species (130). The vectorial competence and role of ticks in WNV transmission remains unclear as limited work has been done on the topic (13). Further research in this area could greatly advance our understanding of WNV transmission dynamics and mechanisms of persistence.

## Conclusion

The burden of WNV in Africa remains underrecognized with fragmented data and surveillance systems. The same is true for several other arboviruses. Disease and genomic surveillance systems are mostly lacking or are poorly resourced, and do not capture the true scope of WNV disease burden on the continent. In several countries, there appears to be a lack of clinical and scientific regard for the disease which prevents effective public health interventions and perpetuates the underestimation of its impact. Addressing this requires a comprehensive and integrated One Health surveillance approach. However, building such a system is complex and riddled with challenges, including resource limitations, complex networks, and the need for multidisciplinary collaboration. We do not know the structure of the ideal WNV surveillance framework for each African country or region, but our study provides recommendations starting with affordable point-of-care diagnostics and targeted genomic surveillance in high-priority regions. While the path forward may be uncertain, our analysis offers actionable starting points to begin addressing these gaps.

## Data sharing

The data and scripts used in this analysis are available at https://github.com/CERI-KRISP/WNV_Genomic_Surveillance_Review_Africa.

## Acknowledgements

We acknowledge the Rockefeller Foundation (HTH 017), the National Institute of Health USA (U01 AI151698) for the United World Antiviral Research Network, and the INFORM Africa project through the Institute of Human Virology Nigeria (U54 TW012041), Global Health EDCTP3 Joint Undertaking and its members and the Bill & Melinda Gates Foundation (101103171), European Union (EU) Horizon Europe Research and Innovation Programme (101046041), the Health Emergency Preparedness and Response Umbrella Program, managed by the World Bank Group (TF0B8412), the Medical Research Foundation (MRF-RG-ICCH-2022-100069), and the Wellcome Trust (228186/Z/23/Z).

