## Supplementary information for "Identifying genomic surveillance gaps in Africa for the global public health response to West Nile Virus"

**Supplementary information S1:** Table of search terms employed in the literature search.

| Search | Terms |
| --- | --- |
| #1 | "West Nile virus"[Mesh] OR "West Nile Fever"[Mesh] OR “West nile virus”[tiab] OR WNV[tiab] OR “West nile fever”[tiab] OR “West nile encephalitis”[tiab] OR “West nile meningitis”[tiab] OR “West nile poliomyelitis”[tiab]  37,502 search results |
| #2 | Africa[tiab] OR Algeria[tiab] OR Angola[tiab] OR Benin[tiab] OR Botswana[tiab] OR "Burkina Faso"[tiab] OR Burundi[tiab] OR "Cabo Verde"[tiab] OR Cameroon[tiab] OR "Canary Islands"[tiab] OR "Cape Verde"[tiab] OR "Central African Republic"[tiab] OR Chad[tiab] OR Comoros[tiab] OR Congo[tiab] OR "Democratic Republic of Congo"[tiab] OR DRC[tiab] OR Djibouti[tiab] OR Egypt[tiab] OR "Equatorial Guinea"[tiab] OR Eritrea[tiab] OR Eswatini[tiab] OR Ethiopia[tiab] OR Gabon[tiab] OR Gambia[tiab] OR Ghana[tiab] OR Guinea[tiab] OR "Guinea Bissau"[tiab] OR "Ivory Coast"[tiab] OR "Cote d'Ivoire"[tiab] OR Jamahiriya[tiab] OR Kenya[tiab] OR Lesotho[tiab] OR Liberia[tiab] OR Libya[tiab] OR Madagascar[tiab] OR Malawi[tiab] OR Mali[tiab] OR Mauritania[tiab] OR Mauritius[tiab] OR Mayotte[tiab] OR Morocco[tiab] OR Mozambique[tiab] OR Namibia[tiab] OR Niger OR Nigeria OR Principe OR Reunion OR Rwanda OR "Sao Tome" OR Senegal[tiab] OR Seychelles[tiab] OR "Sierra Leone"[tiab] OR Somalia[tiab] OR "South Africa"[tiab] OR “Southern Provinces”[tiab] OR "St Helena"[tiab] OR Sudan[tiab] OR Swaziland[tiab] OR Tanzania[tiab] OR Togo[tiab] OR Tunisia[tiab] OR Uganda[tiab] OR "Western Sahara"[tiab] OR Zaire[tiab] OR Zambia[tiab] OR Zimbabwe[tiab]  1,254,281 search results |
| #3 | "Genetic Variation" [Mesh] OR "Genome, Viral" [Mesh] OR Genomics [Mesh] OR Phylogeny [Mesh] OR Metagenome [Mesh] OR Lineage [tiab] OR Strain [tiab] OR Genotype [tiab] OR "Molecular epidemiology" [Mesh] OR "Nucleic acids" [Mesh] OR Genetic [tiab] OR Phylogenetic* [tiab] OR Phylogeo* [tiab] OR Gene [tiab]  4,538,579 search results |
| #4 | NOT Review [publication type] |

**Supplementary information S2:** Publication identification, screening, and inclusion results for literature review according to PRISMA criteria.

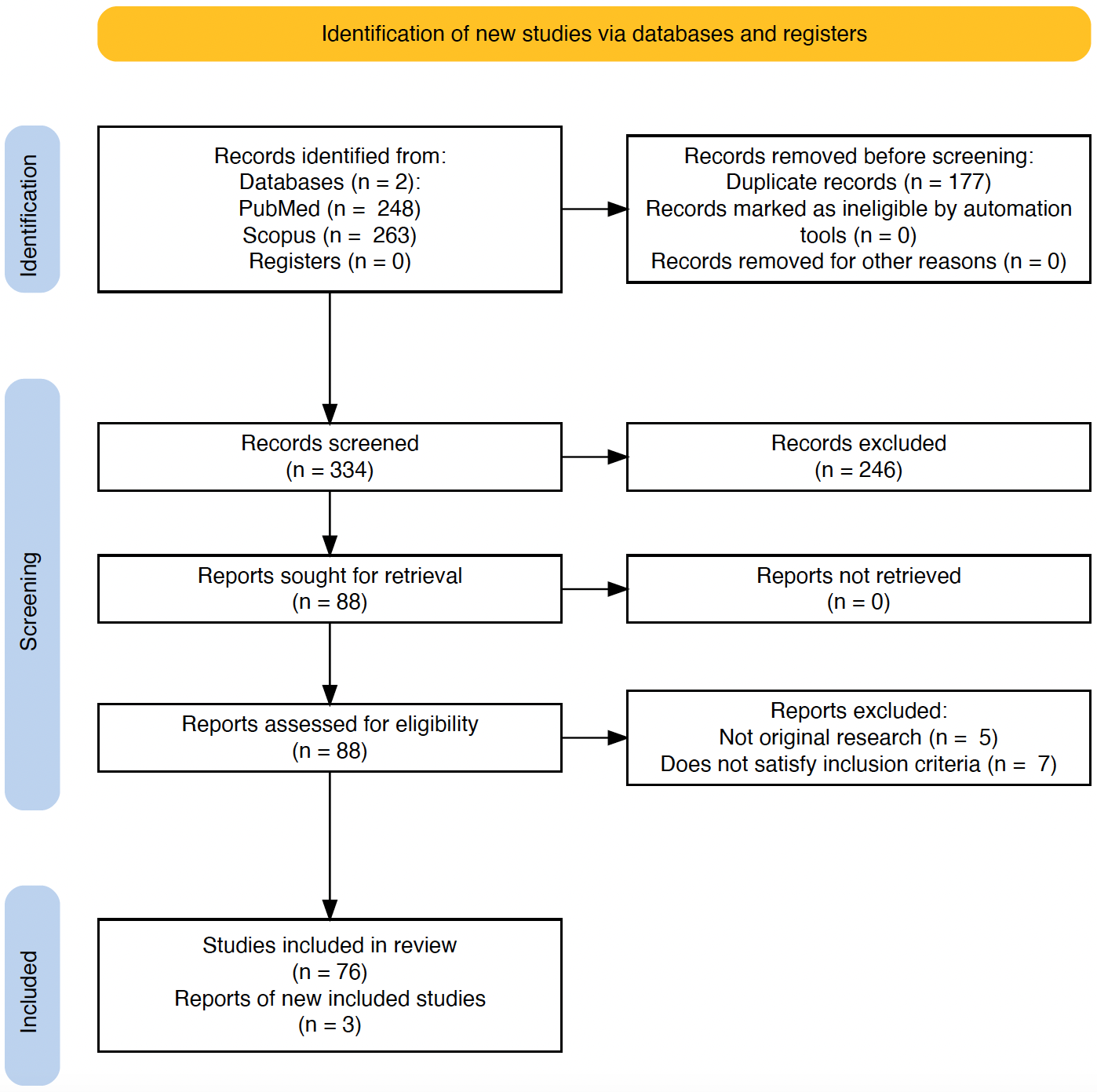

**Supplementary information S3:** Large treemap depicts the number of West Nile virus sequences retrieved from the literature review (n=258) of which 26 are unpublished. The smaller treemap shows the composition of accessioned sequences that were not captured by the literature review (n=58) and that 40 sequences are not linked to publications. Table lists the accession numbers in each of these categories.

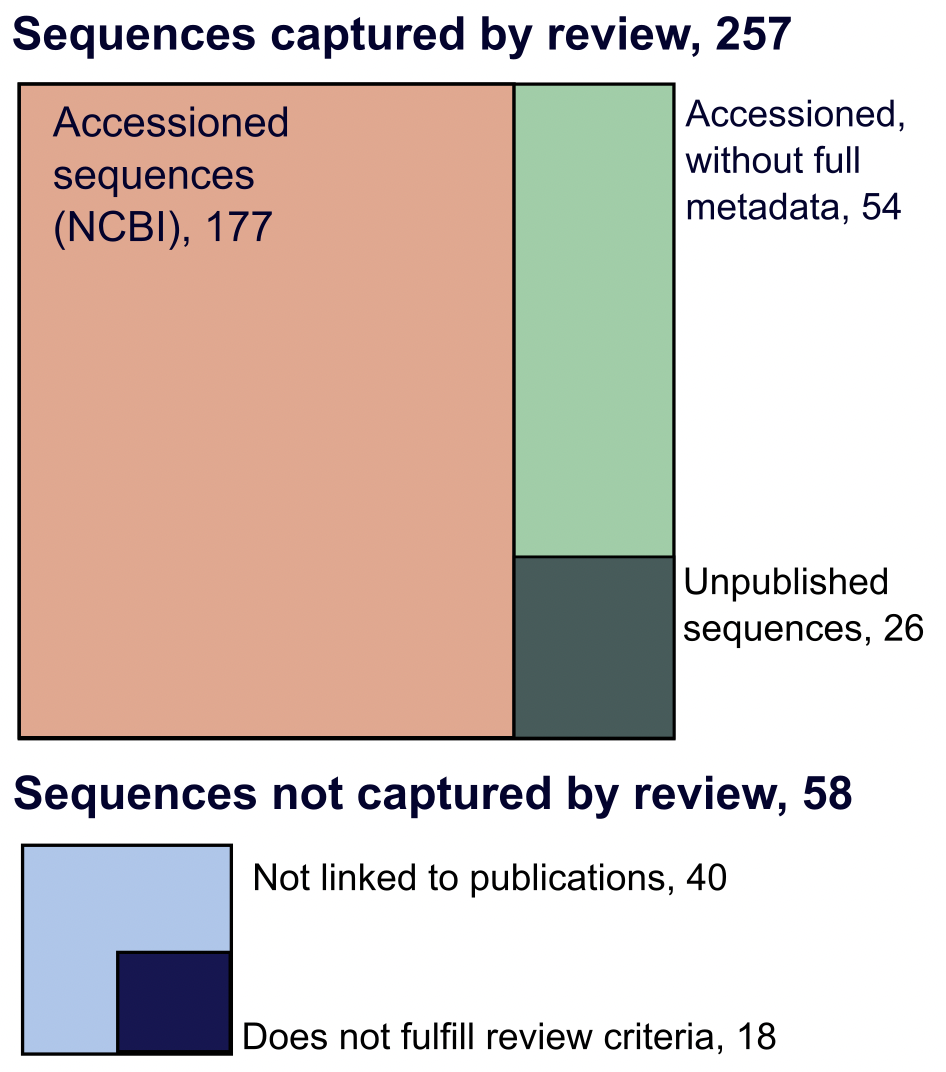

| **Captured by review** |  |
| --- | --- |
| Accessioned sequences | AF146082,AY701412,AY701413,EF429197,EF429198,EF429199,EF429200,FJ464377,  FJ464378,FJ464379,FJ464380,FJ464381,GQ851606,GQ851607,GQ851608,HQ594469,  HQ594470,JN393308,JX974605,KC243146,KP099553,KP099554,KP099555,KP099556,  KP099557,KP099558,KP099559,KP099560,KP099561,KX181348,KX181349,KX181350,  KX181351,KX181352,KX181353,KX181354,KX189174,KX189175,KX189176,KY176717,  KY176718,KY176719,KY176720,KY176721,KY176722,KY176723,KY176724,KY176725,  KY176726,KY176727,KY176728,KY176729,KY176730,KY176731,KY176732,KY176733,  KY176734,KY176735,KY176736,KY703854,KY703855,KY703856,LC318700,LC489409,  LC518897,MF371349,MF371350,MF371351,MF371352,MF371353,MF371354,MF371355,  MF371356,MF371357,MF371358,MF371359,MF371360,MN270988,MN270989,MN270990,  MT055864,MT055865,MT055866,MT055867,MT055868,MT055869,MT055870,MT055871,  MT055872,MT055873,MT055874,MT055875,MT055876,MT055877,MT055878,MT055879,  MT055880,MT055881,MT055882,MT055883,MT055884,MT055885,MT055886,MT132377,  MT132378,MT132379,MT132380,MT132381,MT132382,MT132383,MT132384,MT132385,  MT132386,MT132387,MT132388,MT132389,MW383507,MW383508,MW383509,MW436414,  MW436415,MW436416,MW436417,MW436418,MW436419,MW436420,OL411950,OL411951,  OL411952,OL411953,OL411954,OL411955,OL411956,OL411957,OL411958,OL411959,  OL411960,OL411963,OL411964,OL411965,OL411966,OL411967,OL790166,OL790167,  OM728607,ON813210,ON813211,ON813212,ON813213,ON813214,ON813215,ON813216,  ON813217,ON813218,ON813219,FJ464376,OP846971,OP846972,OP846973,OP846974,  OP846975,OP846976,OP846977,OP846978,OP846979,OP846980,OP846981,OP846982,  OP870453,OP870454,OP870455,OP870456,OP870457,OP870458,OP870459,OP870460,  OP870461 |
| Accessioned but  without full metadata | AY268133,AY262283,AF001556-AF001574,MN057643,PP445046,AF514918-AF514946,  AF260968,M12294 |
| **Not captured by review** |  |
| Not linked to publictions | FJ464382,DQ055173,DQ318019,DQ318020,EU081844,HQ594467,HQ594468,JN226820,  JN226821,JN226822,JN226823,JN226824,JN226825,JN226826,JN226827,JN226828,  JN226829,JN226830,JN226831,JN226832,KM052152,KU978767,KY523178,OL790150,  OL790151,OL790152,OL790153,OL790154,OL790155,OL790156,OL790157,OL790158,  OL790159,OL790160,OL790161,OL790162,OL790163,OL790164,OL790165,OP345226 |
| Does not fulfill review criteria | AF205880,AF205884,AF394221,AY603654,AY839588,AY839589,AY839590,DQ176636,  EU068667,EU074034,HM147822,HM147823,HM147824,KJ131500,KJ131501,KJ131502,  KJ131503,KJ131504 |

**Supplementary information S4:** Results of West Nile virus Lineage 1A phylogenetic tree root to tip regression

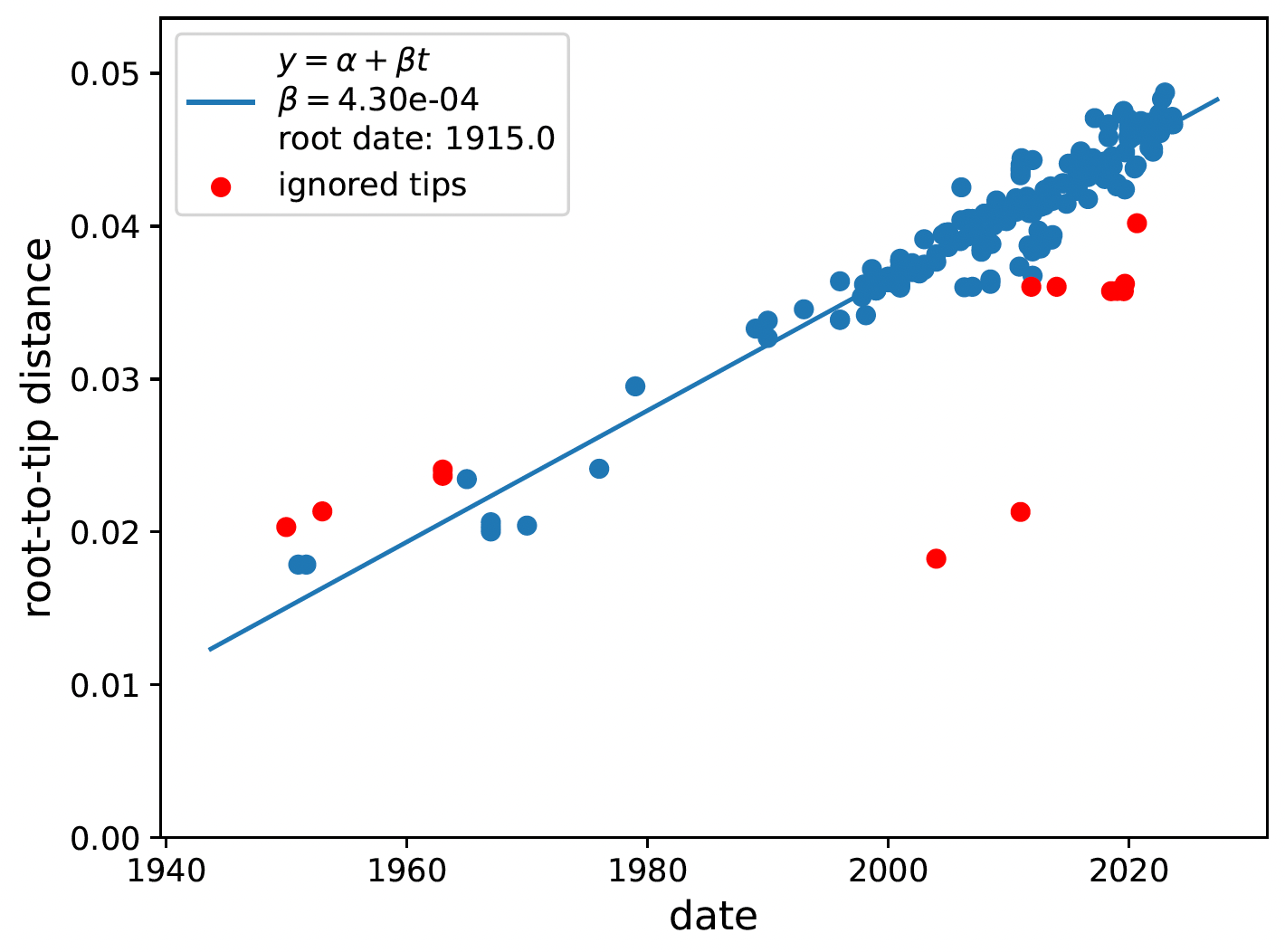

**Supplementary information S5:** Results of West Nile virus Lineage 2 phylogenetic tree root to tip regression

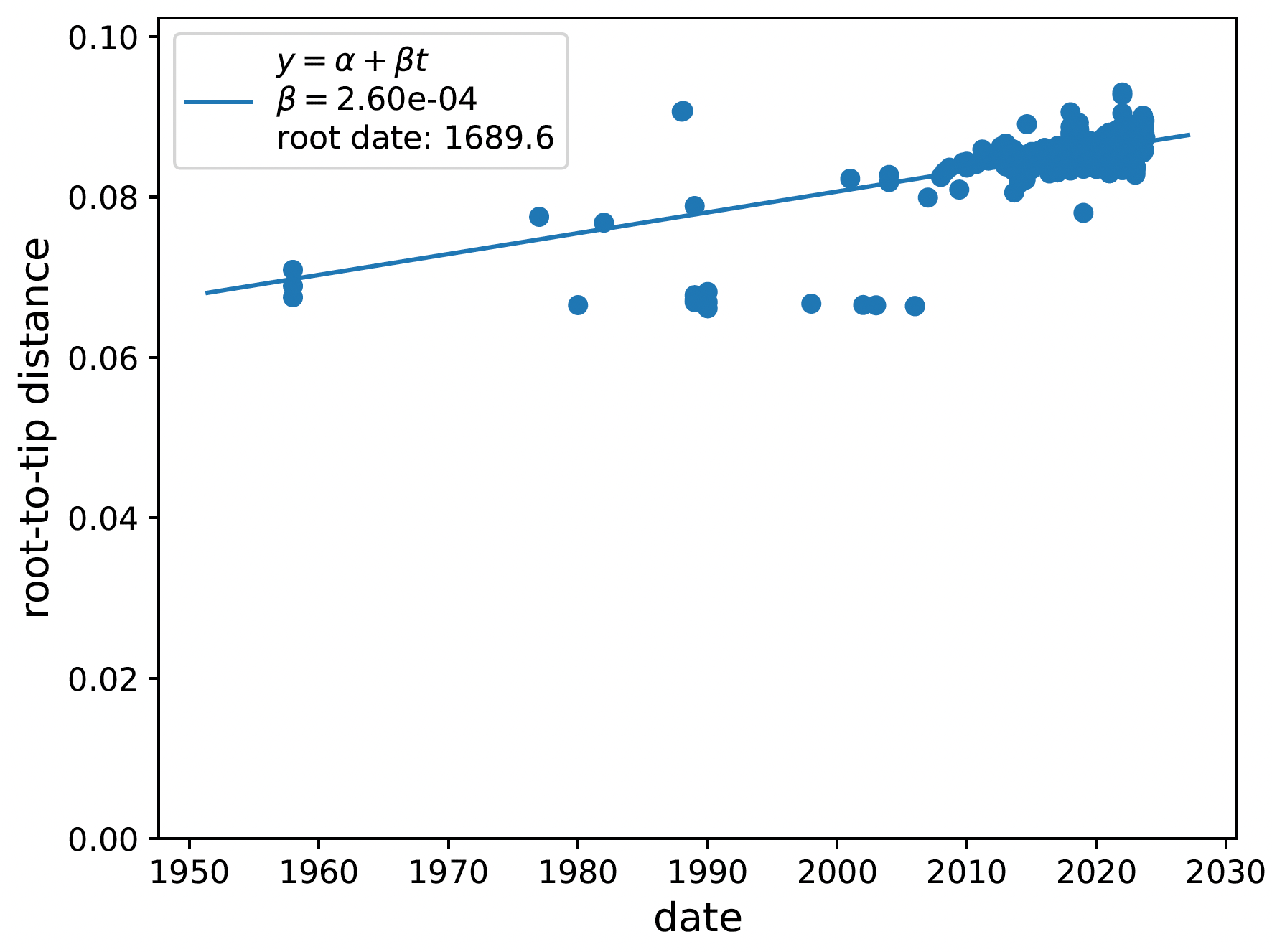

**Supplementary information S6:** A) Point locations (blue) of samples screened by studies with polymerase chain reaction (PCR) methods. Dark grey colouring portrays the countries from which samples were collected. Gabon is zoomed in for resolution of sample locations in the side panel; B) Point locations (gold) of samples collected for genomic sequencing. Tunisia and South Africa are zoomed in on the side panels

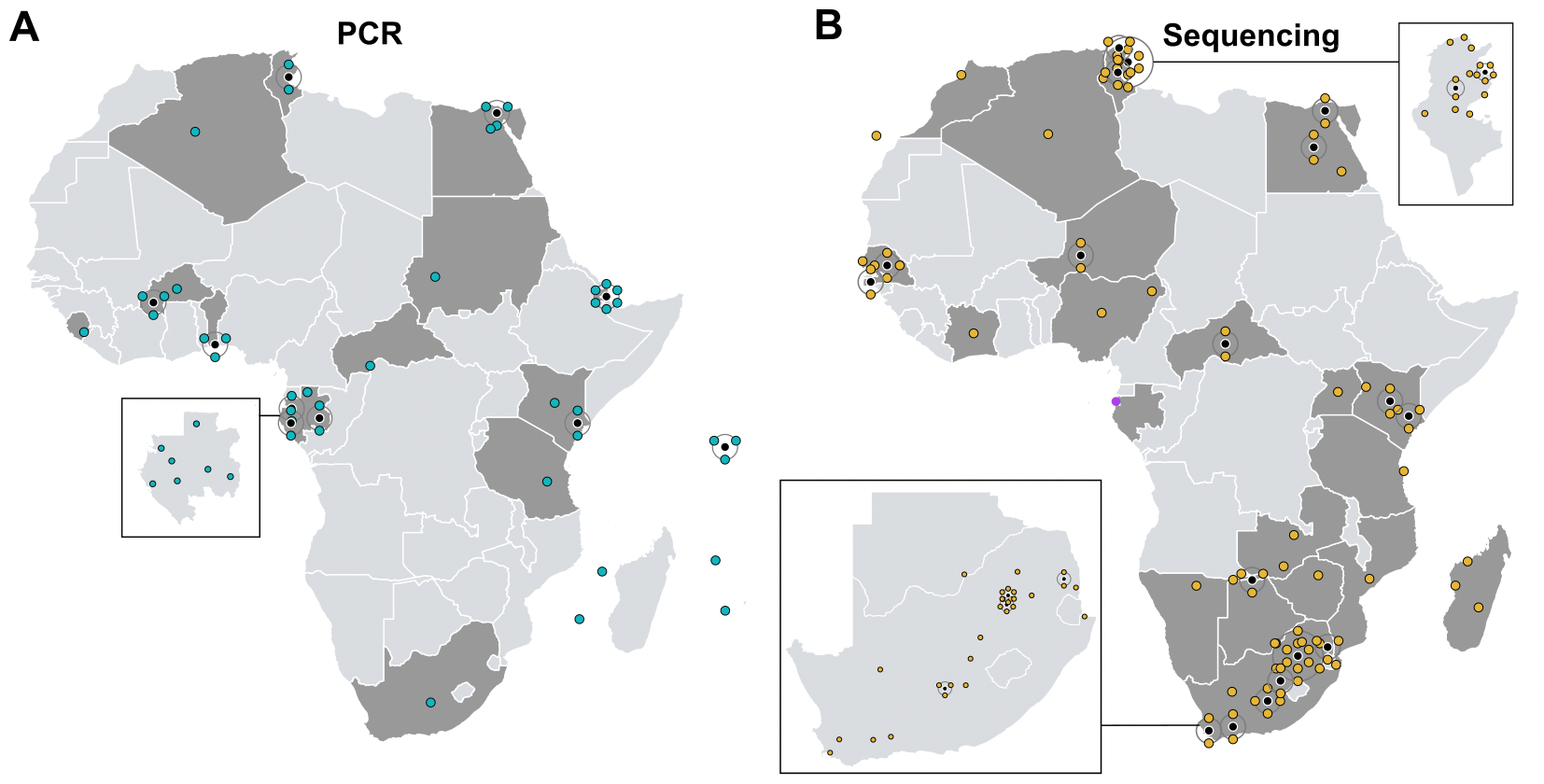

**Supplementary information S7:** Temporal distribution of genomic publications included in the review (grey squares) per country as well as the collection date and the number of publicly available sequences shown by orange circles.

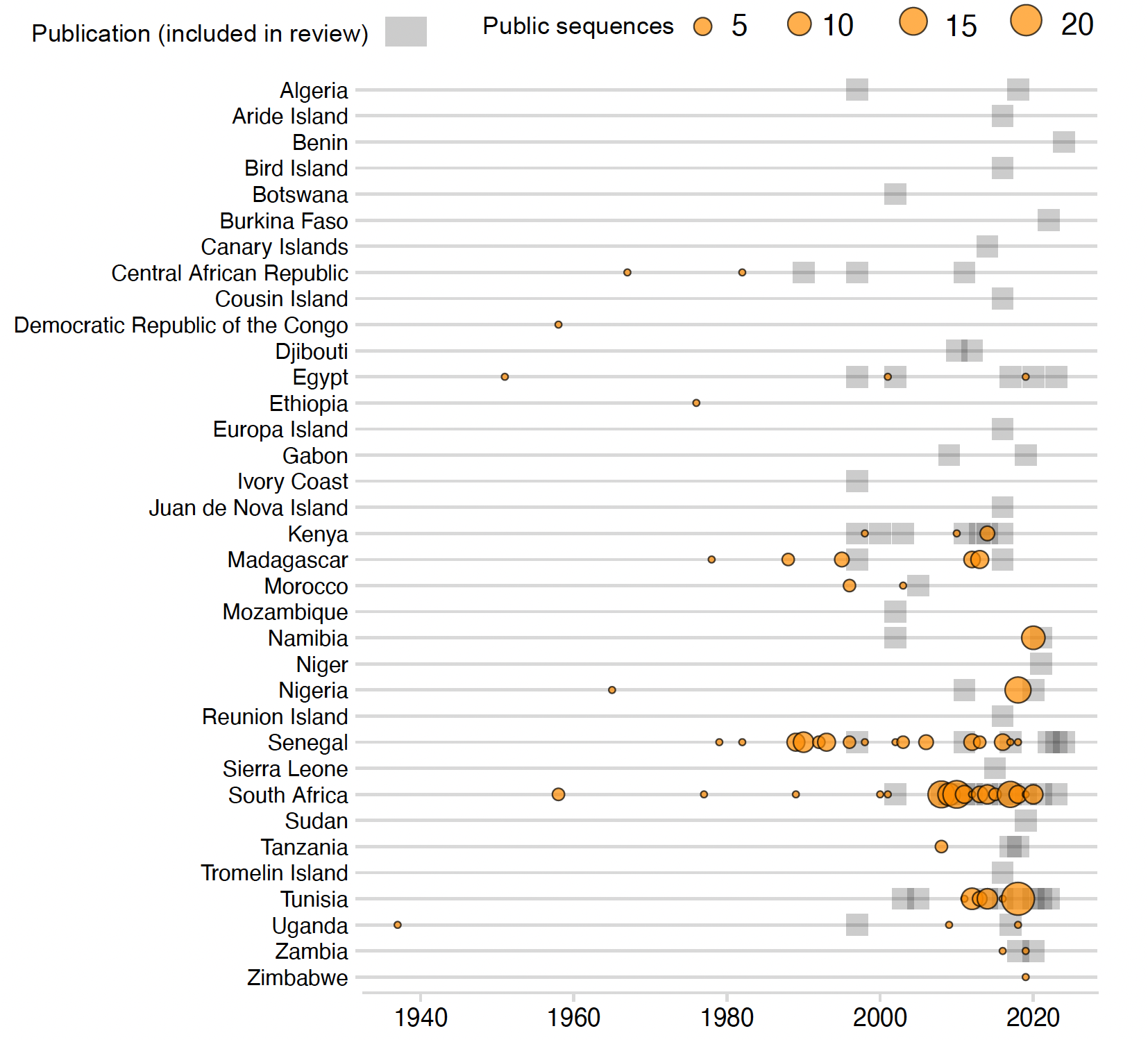

**Supplementary information S8:** For studies that produced African genomic data, the large pie chart depicts the proportion of publications that performed molecular laboratory protocols internally (i.e., within the African country where samples were collected) or externally (i.e., sent to another laboratory). The smaller pie chart displays the locations for studies that employed external laboratory services.

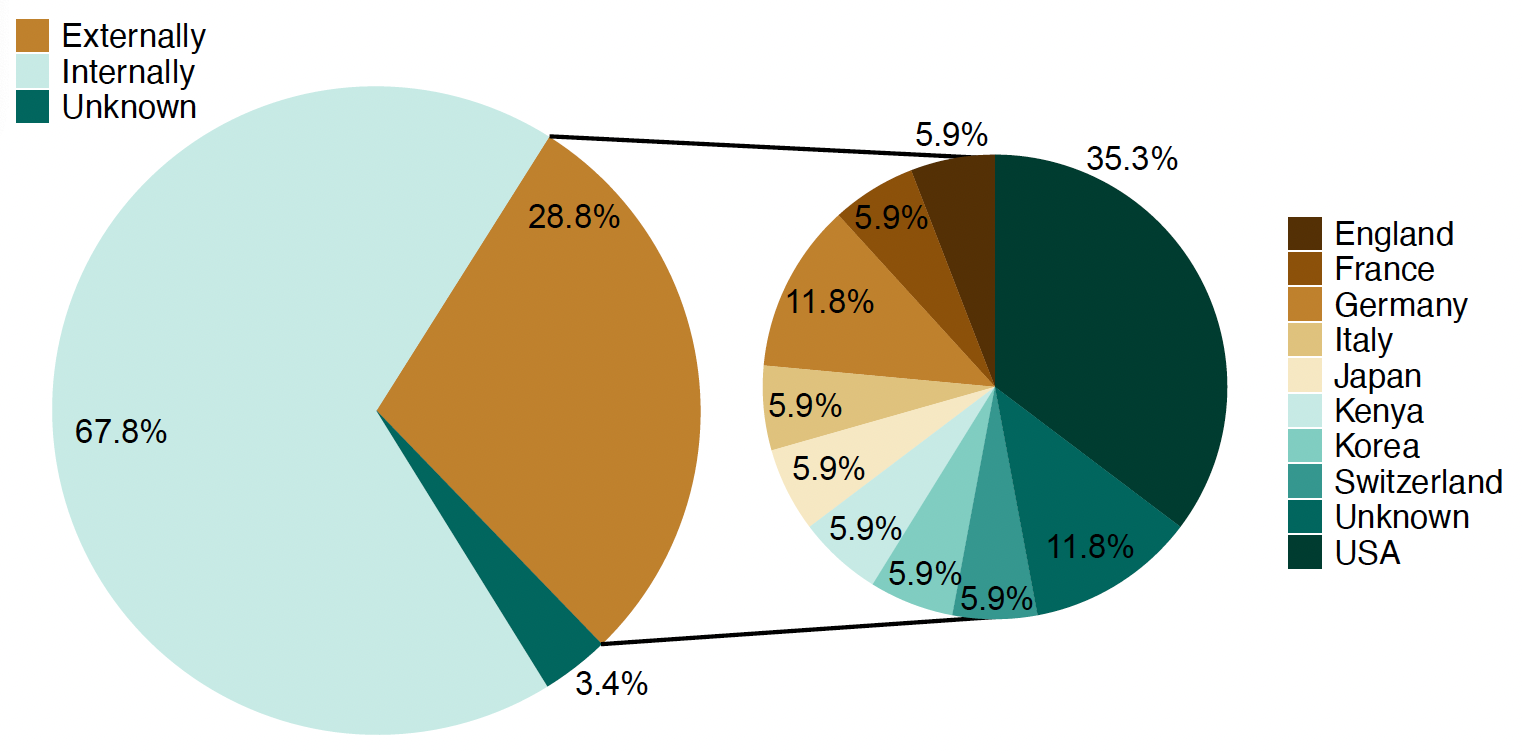

**Supplementary information S9:** Number of publications that employed molecular methods of polymerase chain reaction (PCR) (solely) or sequenced genomic regions per study type.

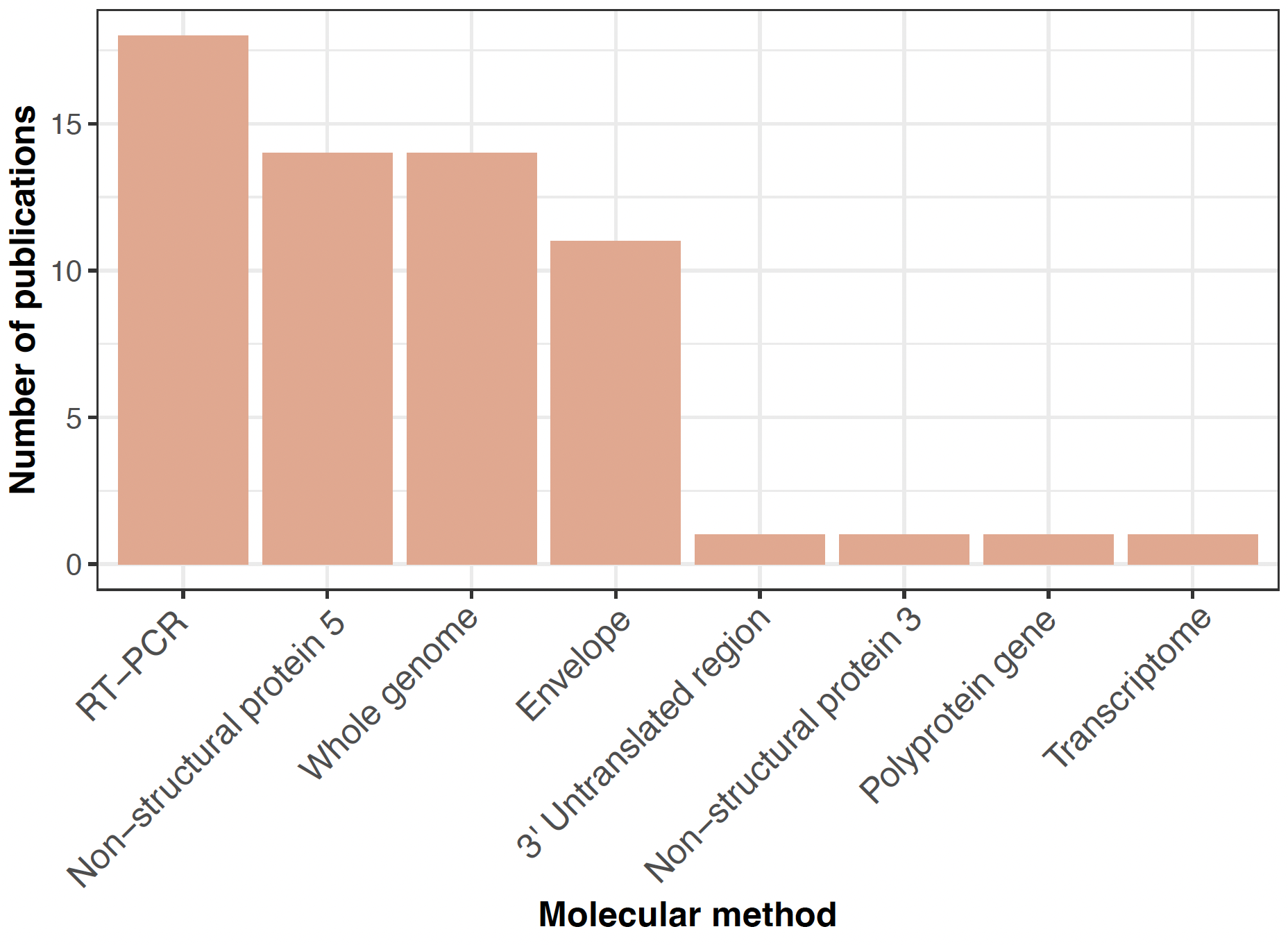

**Supplementary information S10:** Temporal progression of the use of different sequencing methods summed by publications per year.

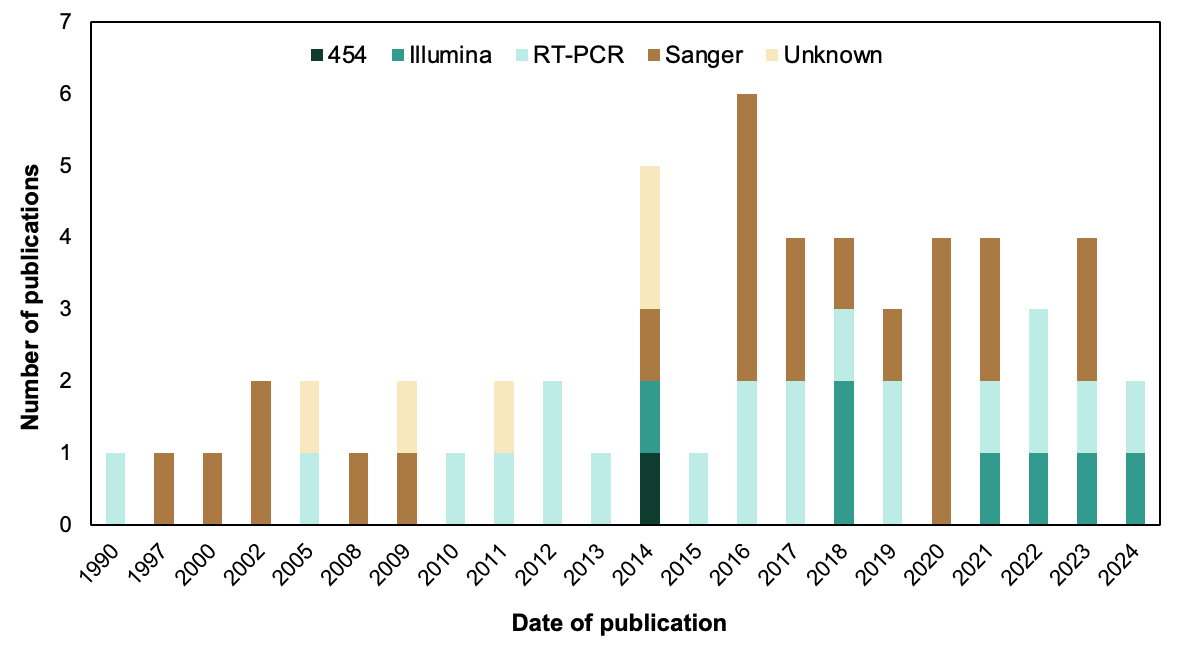
